# Gli3 regulates vomeronasal neurogenesis, olfactory ensheathing cell formation and GnRH-1 neuronal migration

**DOI:** 10.1101/643155

**Authors:** Ed Zandro M. Taroc, Ankana Naik, Jennifer M. Lin, Nicolas B. Peterson, David L. Keefe, Elizabet Genis, Gabriele Fuchs, Ravikumar Balasubramanian, Paolo E. Forni

## Abstract

During mammalian development, gonadotropin-releasing-hormone-1 neurons (GnRH-1ns) migrate from the developing vomeronasal organ (VNO) into the brain asserting control of pubertal onset and fertility. Recent data suggest that correct development of the olfactory ensheathing cells (OEC) is imperative for normal GnRH-1 neuronal migration. However, the full ensemble of molecular pathways that regulate OEC development remains to be fully deciphered. Loss-of-function of the transcription factor Gli3 is known to disrupt olfactory development, however, if Gli3 plays a role in GnRH-1 neuronal development is unclear. By analyzing Gli3 extra-toe mutants (Gli3^Xt/Xt^), we found that Gli3 loss-of-function compromises the onset of achaete-scute family bHLH transcription factor 1 (Ascl-1) positive vomeronasal progenitors and the formation of OEC in the nasal mucosa. Surprisingly, GnRH-1 neurogenesis was intact in Gli3^Xt/Xt^ mice but they displayed significant defects in GnRH-1 neuronal migration. In contrast, Ascl-1^null^ mutants showed reduced neurogenesis for both vomeronasal and GnRH-1ns but less severe defects in OEC development. These observations suggest that Gli3 is critical for OEC development in the nasal mucosa and subsequent GnRH-1 neuronal migration. However, the non-overlapping phenotypes between Ascl-1 and Gli3 mutants indicate that Ascl-1, while crucial for GnRH-1 neurogenesis, is not required for normal OEC development. Since Kallmann syndrome (KS) is characterized by abnormal GnRH migration, we examined whole exome sequencing data from KS subjects. We identified and validated a *GLI3* loss-of-function variant in a KS individual. These findings provide new insights into GnRH-1 and OECs development and demonstrate that human *GLI3* mutations contribute to KS etiology.

**Significance statement:** The transcription factor Gli3 is necessary for correct development of the olfactory system. However, if Gli3 plays a role in controlling GnRH-1 neuronal development has not been addressed. We found that Gli3 loss-of-function compromises the onset of Ascl1+ vomeronasal progenitors, formation of olfactory ensheathing cells in the nasal mucosa and impairs GnRH-1 neuronal migration to the brain. By analyzing Ascl1 null mutants we dissociated the neurogenic defects observed in Gli3 mutants from lack of olfactory ensheathing cells in the nasal mucosa, moreover, we discovered that Ascl1 is necessary for GnRH-1 ontogeny. Analyzing human whole exome sequencing data, we identified a *GLI3* loss-of-function variant in a KS individual. Our data suggest that *GLI3* is a candidate gene contributing to KS etiology.

## Introduction

The olfactory placode gives rise to multiple cell types, including migratory/pioneer neurons, terminal nerve cells, gonadotropin releasing hormone-1 (GnRH-1) neurons (GnRH-1ns), olfactory, and vomeronasal sensory neurons (VSNs) (Wray et al., 1989; Miller et al., 2010; Casoni et al., 2016). GnRH-1ns play a central role in controlling the reproductive axis of vertebrates. During embryonic development GnRH-1ns migrate along the terminal nerve (Schwanzel-Fukuda and Pfaff, 1989; Casoni et al., 2016; Taroc et al., 2017) from the vomeronasal organ into the hypothalamus. Once in the brain the GnRH-1ns control gonadotropin release from the pituitary gland. Defects in neurogenesis and/or migration of GnRH-1ns can lead to Kallmann syndrome (KS), a genetic disorder characterized by pubertal failure due to congenital hypogonadotropic hypogonadism (HH) and anosmia (impaired sense of smell) (Pingault et al., 2013; Forni and Wray, 2015).

In the last three decades, various causal genes for KS have been identified and disruption of these genetic pathways lead to KS through aberrant development of olfactory placode derivatives and/or impaired migration of GnRHns. Notably, recent studies show that the ability of the GnRH-1ns to migrate from the olfactory pit to the brain is also critically dependent on the correct development of a specialized neural crest derived glial cells termed “olfactory ensheathing cells” (Barraud et al., 2013; Pingault et al., 2013). In keeping with this notion, of the known KS genes, two genes, SOX10 and *ANOS1* have been recently identified as key factors controlling OECs development (Hu et al., 2019). Thus, understanding the genetic pathways governing OEC development could provide more mechanistic clues into the basis of aberrant GnRH neuronal migration and therefore, KS.

Gli3, together with Gli1 and Gli2, are key transcription factors involved in the sonic hedgehog (Shh) signaling. In the absence of Shh signaling, Gli3 and Gli2 act as transcriptional repressors. However, in the presence of Shh, Gli1,2 and Gli3 function as transcriptional activators (Sasaki et al., 1999; Niewiadomski et al., 2014). A spontaneous murine model of the *Gli3* gene, Gli3^pdn/pdn^ has previously implicated a potential role for Gli3 in GnRH neuronal migration. While the hypomorphic Gli3^pdn/pdn^ model shows a delayed, but not missing, GnRH-1 neuronal migration (Naruse et al., 1994), the more severe Gli3 null mutants Gli3 extratoe (Gli3^Xt/Xt^) fail to develop a functional olfactory system (Keino et al., 1994; Balmer and LaMantia, 2004; Besse et al., 2011). However, GnRH1 neuronal development and migration in Gli3^Xt/Xt^ has not been analyzed.

A candidate gene study in humans has revealed missense variants in the *GLI3* gene in HH (Quaynor et al., 2016) but no functional validation has been performed to ascertain the causality of these variants. Furthermore, *GLI3* mutations in humans have been reported in non-syndromic forms of polydactyly and syndromic forms of polydactyly of which a subset of patients display neonatal hypogonadism (micropenis and undescended testes) (Johnston et al., 2010). Despite these observations, the precise role of Gli3 in GnRH neuronal biology remains to be fully elucidated.

By analyzing *Gli3*^Xt/Xt^ mutants across the key early developmental stages of embryonic olfactory and GnRH neuronal development, we found that *Gli3* loss-of-function compromises the onset of Ascl1+ vomeronasal progenitors and disrupts the formation of OECs in the nasal mucosa. Notably, in Gli3^Xt/Xt^, although GnRH-1 neurogenesis was preserved as in controls, the GnRHns were unable to migrate and populate the hypothalamus. To further dissect the precise roles of Gli3 and Ascl1, we also analyzed *Ascl1* null mutants and found significant reduction for both vomeronasal and GnRH-1ns but less severe defects in OECs’ development than in Gli3^xt/xt^, suggesting the OEC development is critically dependent on *Gli3* rather than *Ascl1*. By whole exome sequencing of human KS patients, we identified and functionally validated the first *GLI3* loss-of-function variant in a KS individual. These findings provide new insights into GnRH-1 and OECs development and demonstrate that human *GLI3* mutations contributes to KS etiology.

## Material and Methods

### Animals

Gli3^Xt^ (Schimmang et al., 1992) and Ascl-1^tm1.1(Cre/ERT2)Jeo^/J mice (Kim et al., 2011) on C57BL/6J background were purchased from (Jackson Laboratories, Bar Harbor, ME). Both colonies were maintained on C57BL/6J. The genotypes of Gli3^Xt^ mice were established by PCR analysis using the following primers: Gli3-C3F: GGCCCA AACATCTACCAACACATAG, Gli3-C3R: GTTGGCTGCTGCATGAAGACTGAC; Gli3-XtJ580F: TACCCCAGCAGGAGACTCAGATTAG; Gli3-XtJ580F: AAACCCGTGGCTCAGGACAAG. Ascl-1^CreERT2^ were genotyped following Ascl-1^tm1.1(Cre/ERT2)Jeo^/J protocol available on jax.org website. Amplification products were analyzed by agarose gel electrophoresis. Animals were euthanized using CO2, followed by cervical dislocation. Mutant and wild-type mice of either sex were used. All mouse studies were approved by the University at Albany Institutional Animal Care and Use Committee (IACUC).

### Tissue preparation

Embryos were collected from time-mated dams where the emergence of the copulation plug was taken as E0.5. Collected embryos were immersion-fixed in 3.7% Formaldehyde/PBS at 4°C for 3-5 hours. Postnatal animals were perfused with 3.7% Formaldehyde/PBS. All samples were then cryoprotected in 30% sucrose overnight, then frozen in O.C.T (Tissue-TeK) and stored at −80°C. Samples were cryosectioned using CM3050S Leica cryostat and collected on Superfrost plus slides (VWR) at 12-16μm thickness for immunostainings and 18-25μm for *in situ hybridizations (ISH)*.

### Immunohistochemistry

Primary antibodies and dilutions used in this study were: goat-α-neuropilin-2 (1:3000, R&D Systems), goat-α-neuorpilin-1 (1:400, R&D Systems), mouse-α-ROBO2(1:50, Santa Cruz) chicken-α-peripherin (1:1500, Abcam), rabbit-α-peripherin (1:2000, Millipore), SW rabbit-α-GnRH-1 (1:6000, Susan Wray, NIH), goat-α olfactory marker protein (1:4000, WAKO), mouse -α-GAD67 (1:200, Santa Cruz), goat-α-AP-2ε (1:1000, Santa Cruz), mouse-α-Meis2 (1:500, Santa Cruz), rabbit-α-phospho-Histone-H3 (1:400, Cell Signaling), rat-α-phospho-Histone-H3 (1:500, Abcam), goat-α-Sox10 (1:100, Santa Cruz), goat-α-collagen IV (1:800, Millipore), goat-α-Gli3 (1:200, R&D Systems), rabbit-α-Hes-1 (1:200, Cell Signaling), and mouse-α-Ascl-1 (1:30, Santa Cruz). Antigen retrieval was performed in a citrate buffer prior to incubation with chicken-α-peripherin, rabbit-α-phospho-Histone-H3, rat-α-phospho-Histone-H3, goat-α-AP-2ε, mouse-α-Meis2, goat-α-Gli3, rabbit-α-Hes-1, mouse-α-Ascl-1, goat-α-Sox10, mouse-α-ROBO2, and mouse-α-GAD67 antibodies. For immunoperoxidase staining procedures, slides were processed using standard protocols (Forni et al., 2013) and staining was visualized (Vectastain ABC Kit, Vector) using diaminobenzidine (DAB) in a glucose solution containing glucose oxidase to generate hydrogen peroxide; sections were counterstained with methyl green. For immunofluorescence, species-appropriate secondary antibodies were conjugated with Alexa-488, Alexa-594, or Alexa-568 (Molecular Probes and Jackson Laboratories) as specified in the legends. Sections were counterstained with 4′,6′-diamidino-2-phenylindole (1:3000; Sigma-Aldrich) and coverslips were mounted with Fluoro Gel (Electron Microscopy Services). Confocal microscopy pictures were taken on a Zeiss LSM 710 microscope. Epifluorescence pictures were taken on a Leica DM4000 B LED fluorescence microscope equipped with a Leica DFC310 FX camera. Images were further analyzed using FIJ/ImageJ software. Each staining was replicated on at least three different animals for each genotype

### In situ hybridization

Digoxigenin-labeled cRNA probes were prepared by in vitro transcription (DIG RNA labeling kit; Roche Diagnostics) from the following templates: Semaphorins 3A (Kagoshima and Ito, 2001), Gli3 riboprobe RP_080717_03_E04 (Forward primer GCAGAATTATTCCGGTCAGTTC; Reverse primer TCAGTCTTTGTGTTTGTGGTCC). In situ hybridization was performed as described (Lin et al., 2018b) and visualized by immunostaining with an alkaline phosphatase conjugated anti-DIG (1:1000), and NBT/BCIP developer solution (Roche Diagnostics). Sections were then counter-immunostained with antibodies against both chicken-α-peripherin, and SW rabbit-α-GnRH-1, as described above for immunofluorescence.

### Cell quantifications

Quantification of GnRH-1 neuronal distribution in mutant and control animals was performed in the nasal region (VNO, axonal tracks surrounding the olfactory pits), forebrain junction and brain (all the cells that accessed the olfactory bulb and were distributed within the forebrain). Number of cells were calculated for each animal as the average of the counted cells per series multiplied by the number of series cut per animal. Differences at each interval between genotypes was assessed by unpaired t-Test.

Number of vomeronasal apical and basal neurons was counted after AP-2ε and Meis2 immunostaining, as cells/section on an average of 4 sections/embryo. Progenitors cells were counted after immunostaining for Phospho-Histone H3, Ascl-1, and Hes-1 and were quantified as described above for vomeronasal cells. Sox10 positive cells were quantified in a 334,500 μm^2^ field area for an average of 4 VNO sections per animal, only Sox10 positive cells proximal to the VNO were counted. Frequency of Sox10 cells on vomeronasal bundles was calculated by counting the number of Sox10 I.R. cells/μm fiber length. Means ± SEs were calculated on at least three animals per genotype, *t* test was used to assess differences between groups.

### Densitometric Analysis of the Median Eminence

Brains of 5-month-old male Gli3^Xt^ WT and heterozygous mice (n=3;3) were cryosectioned coronally at 20μm. Three series for each animal were stained for GnRH in DAB. Brightfield images of comparable sections for each series were taken for quantification. Images were processed with color deconvolution and the mean gray value was measured for a standardized ROI to calculate the average optical density for each animal.

### Site directed mutagenesis

The GLI-3 bs-2 plasmid carrying the human GLI3 (Kinzler et al., 1988) was purchased from Addgene. F387F*fs mutation was generated in the vector pCDNA3 hGli3 using Q5 site directed mutagenesis kit (NEB) using primers Gli3 F387Ffs* Fwd CCAACACAGAGGCCTATTCC and Gli3 F387Ffs* Rev AAAGTTGGGGCAGGGTGG following the manufacturer’s instruction. Mutation was validated by sequencing prior to use in experiments.

### Transfection and luciferase reporter assay

1×10^4^ HeLa cells (ATCC CCL2) were seeded into a 96-well plate 24 hours before transfections. 100 ng of either reporter plasmid (8 × 3 GLI-BS Luc or 8 × 3 Mut GLI-BS Luc) and 100 ng of the effector plasmid (hGli3 WT or F387F*fs) and 20 ng of pRL-SV40 were co-transfected using Lipofectamine 3000 (ThermoFisher). pRL-SV40 coding for *Renilla* luciferase was used as a transfection control and for normalization. Cells were harvested 48 hours post transfection and Firefly and *Renilla* luciferase intensities were measured using Glomax 96 microplate luminometer (Promega) and dual-Luciferase Reporter Assay System (Promega).

### Human phenotyping and genotyping

#### WES analysis of KS individuals for *GLI3* loss-of-function mutations

A total of 632 KS subjects at the Harvard Reproductive Endocrine Sciences Center were examined for *GLI3* loss-of-function mutations in this study. KS was defined by: (i) absent or incomplete puberty by age 18y; (ii) serum testosterone <100 ng/dL in men or estradiol<20 pg/mL in women in the face of low or normal levels of serum gonadotropins; (iii) otherwise normal anterior pituitary function; (iv) normal serum ferritin concentrations; and (v) normal MRI of the hypothalamic-pituitary region (Balasubramanian and Crowley, 2011); and (v) impaired olfaction as determined by the University of Pennsylvania Smell Identification Test (UPSIT) (Lewkowitz-Shpuntoff et al.; Doty, 2007). Clinical charts, biochemical testing (including overnight neuroendocrine testing for LH pulse profiles) and patient questionnaires of KS subjects were reviewed for phenotypic evaluation as described previously (Pitteloud et al., 2002). Whole exome sequencing (WES) data in the KS cohort was performed on peripheral blood-derived DNA using either Nimblegen SeqCap target enrichment kit (Roche) or a custom Illumina capture kit (ICE) and the detailed variant-calling and annotation used have been described previously (Guo et al., 2018). WES data were then queried for *GLI3* (RefSeq: NM_000168.6) loss-of-function variants: defined as those variants resulting in protein alteration in the canonical *GLI3* transcript as determined by Variant Effect Predictor (VEP) (127), resulting in nonsense, and essential splice site mutations; or (b) indels resulting in frame-shift mutations; AND (b) with ethnicity-specific minor allele frequency (MAF) <0.1% minor allele frequency (MAF) in gnomAD database (http://gnomad.broadinstitute.org) (Karczewski et al., 2019). To ascertain oligogenicity, KS proband(s) with *GLI3* loss-of-function variants were also examined for rare sequence variants (RSVs) in other known KS/nIHH genes [*ANOS1* (NM_000216.4), *CHD7* (NM_017780.4), *FEZF1* (NM_001160264.2), *FGF8* (NM_006119.4), *FGFR1* (NM_023110.2), *GNRH1* (NM_000825.3), *GNRHR* (NM_000406.2), *HS6ST1* (NM_004807.3), *KISS1* (NM_002256.4), *KISS1R* (NM_032551.5), *NR0B1* (NM_000475.4), *NSMF* (NM_001130969.1), *PROK2* (NM_001126128.2), *PROKR2* (NM_144773.3), *SEMA3A* (NM_006080.3), *SMCHD1* (NM_015295.2), *SOX2* (NM_003106.4), *TAC3* (NM_013251.4), *TACR3* (NM_001059.3), and *WDR11* (NM_018117.12)]. RSVs in these other HH genes were defined as single nucleotide variants resulting in protein alteration in canonical transcript as determined by Variant Effect Predictor (VEP) (127) resulting in missense, nonsense, and essential splice site mutations; or (b) indels resulting in frame-shift mutations; AND (b) with ethnicity-specific minor allele frequency (MAF) <1% in GnoMAD. Variants identified from above analysis were then confirmed by Sanger sequencing. This study was approved by the Partners Institutional Review Board at the MGH, and informed consent was obtained from all participants.

#### Statistical Analyses

All statistical analyses were carried out using GraphPad Prism7 software. Cell counts were done on serial sections immunostained for GnRH-1 at E13.5 (n=3) and E15.0 (n=4) and visualized under bright field (immunoperoxidase) or epi-fluorescence illumination (20×; Leica DM4000 B LED), according to their anatomical location [*i.e.,* (1) nasal region (VNO, axonal tracks surrounding the olfactory pits, forebrain junction); (2) olfactory bulb/fibrocellular mass; and (3) brain (all the cells that accessed the olfactory bulb and were distributed within the forebrain)]. For each animal, counts were performed on 3 serial series. The average number of cells from these 3 series was then multiplied by the total number of series/animal to compute a value for each animal. These were then averaged ± standard error (SE) among animals of the same age and genotype. Means ± SEs were calculated on at least three animals per genotype. The statistical difference between genotypes and groups were determined using an unpaired student’s t-test. A gene-based burden testing for human *GLI3* variants was performed between the KS and the gnomAD using a Fisher’s-exact test. All data are represented as the mean ±SEM from n≥3 mice per genotype/age for each experiment. Values of *p* < 0.05 were considered to be statistically significant.

## RESULTS

### Gli3 is expressed by apical Hes-1+ progenitor cells in the VNO

Since GnRH-1ns originate in the region of the nasal pit that develops into the vomeronasal organ, we analyzed Gli3 expression throughout GnRH-1ns and VSN development. GnRH-1ns arise between E10.5 and E11.5 and are immunodetectable before vomeronasal neuron formation (Fig.1A,A1,M,P) (Forni et al., 2011a; Forni et al., 2013; Forni and Wray, 2015; Taroc et al., 2017). By E13.5 – E15.5, we observed that the developing VNO was populated by Tfap2e (AP-2ε) positive basal VSNs and Meis2 positive apical VSNs (Fig.1N,Q). However, most GnRH-1ns migrated out of the vomeronasal area by E13.5 (Fig.1B,B1,C,C1).

**Figure 1.**
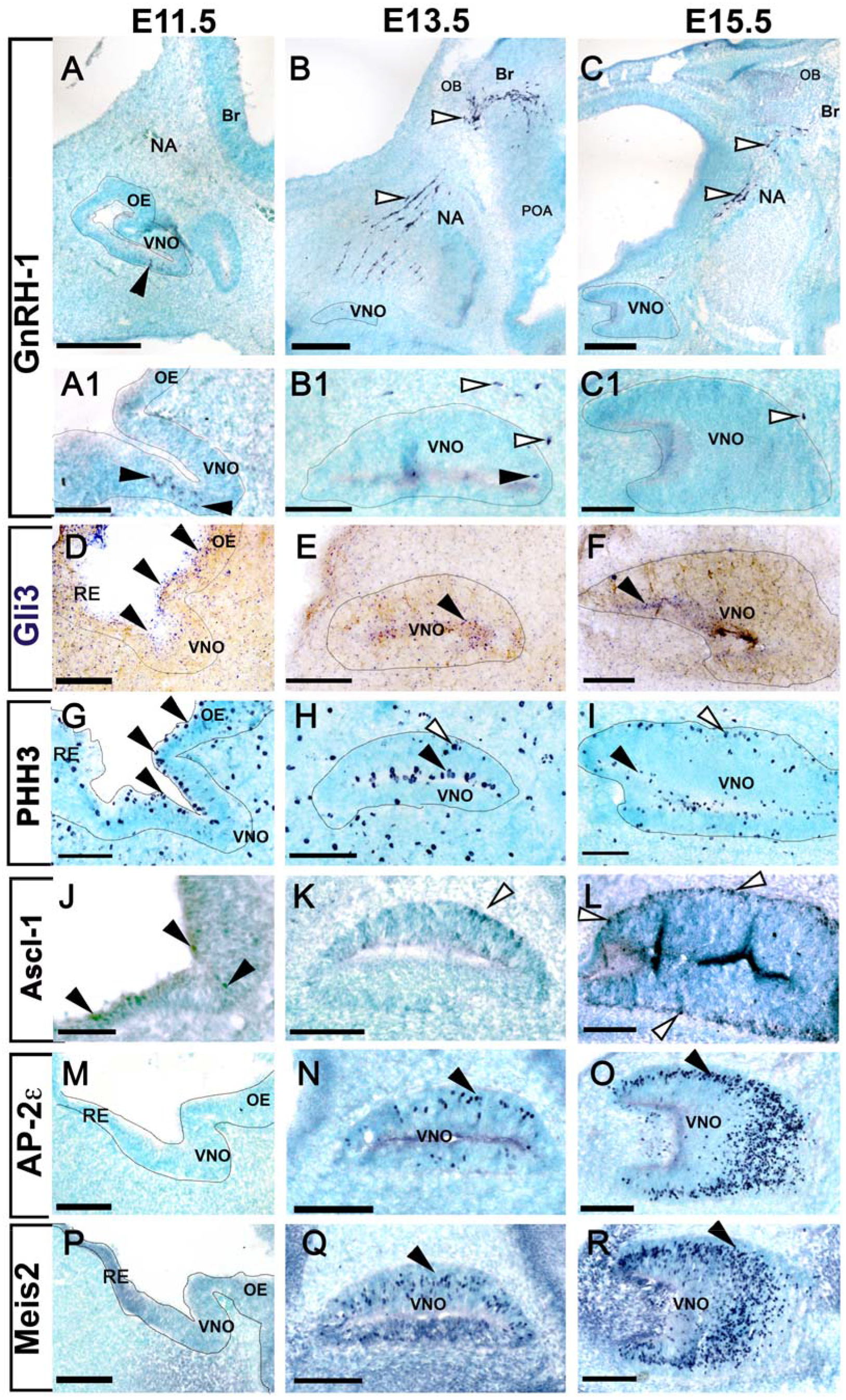
Gli3 is expressed in the developing vomeronasal area. GnRH-1ns can be detected at E11.5 (A,A1) in the ventral portion of the developing putative VNO and emerge from the VNO (white arrows) to the nasal area (NA) migrating towards the brain (Br) at E13.5 (B,B1) and E15.5 (C,C1). D-F Gli3 mRNA expression is detected in cells located in the apical portion of the developing olfactory pit and VNO (black arrows) at all analyzed stages. G-I) PHH3 immunoreactivity highlights dividing cells in the apical (black arrow) and basal (white arrows) regions of the developing VNO. J-L) Immunostaining against Ascl-1 highlights sparse Ascl-1 positive cells at E11.5 (J), neurogenic Ascl-1+ cells were detected in the basal regions of the developing VNO at E13.5 and E15.5. Immunostaining against the basal transcription factor AP-2ε (M-O) and apical transcription factor Meis2 (P-R) reveal the lack of detectable vomeronasal neurons at E11.5, while a growing number of neurons were detected between E13.5 and E15.5. Scale bars A-C: 250 μm, A1,D,G,J,MP: 50 μm, B1,C1,E,F,H,I,K,L,N,O,Q,R: 100 μm

We then performed in-situ hybridization and immunohistochemistry (Data not shown) against Gli3 at embryonic stages E11.5, E13.5, and E15.5 (Fig.1D-F). These experiments revealed that Gli3 mRNA and protein mostly localize in the apical regions of the developing vomeronasal epithelia. The proliferative apical cells in the developing olfactory pit (Cuschieri and Bannister, 1975b) localize Hes-1 positive stem cell/progenitors that give rise, over time, to apical Ascl-1 neurogenic progenitors (Cau et al., 2000). By performing anti phospho-Histone-H3 (PHH3) immunostaining, we only detected mitotic cells in the apical region of the VNO at E11.5. However, at E13.5 and E15.5, we detected mitotic cells along the margins of both the apical and basal regions (Cuschieri and Bannister, 1975a; Cau et al., 2000) (Fig.1G-I). Immunostaining for Hes-1 and Ascl-1 in the developing VNO (Fig.2A-D) showed a progressive enrichment of immune-detectable Ascl1 positive cells over time (Fig.1J-L). We confirmed opposite expression patterns for Hes-1 and Ascl-1 at E15.5, as Hes-1 localized predominantly in the apical portions and Ascl-1 localized in the basal portions of the developing VNO (Fig.2A-D). We confirmed Gli3 immunoreactivity in Hes-1 positive cells and sparse immunoreactivity in the basal regions of the VNO (Fig.2G, H).

**Figure 2.**
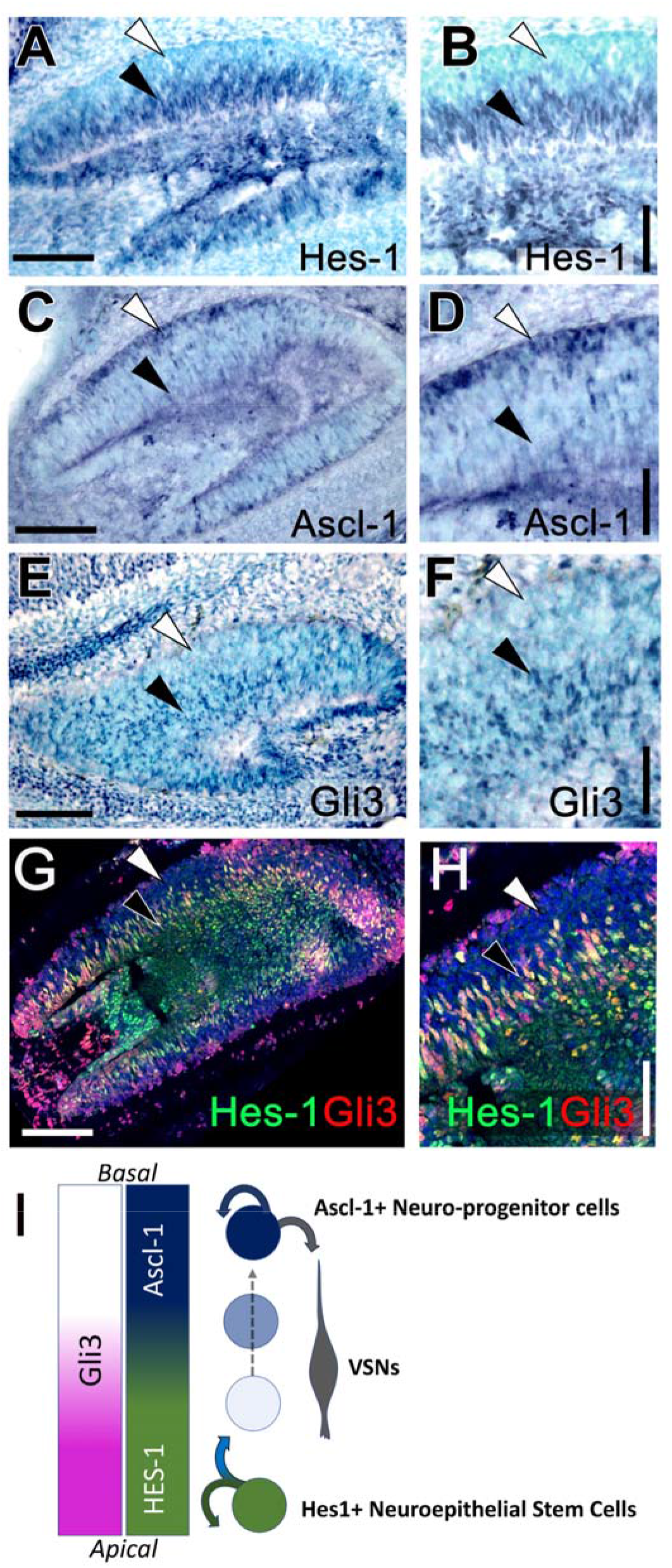
A) Gli3 is expressed in Hes-1, but not in Ascl-1, positive progenitors. E15.5 immunostaining against Hes-1 reveals Hes-1 expression in the cells located in the apical domains of the developing VNO (Black arrow). C,D) Immunostaining against Ascl-1 reveals expression of the neurogenic factor in cells in the basal territories of the developing VNO (White arrow), only background levels were found in the basal regions (Black arrow). E,F) Gli3 immunoreactivity was found in cells in the apical territories of the developing VNO (Black arrow). G-H) Double immunofluorescence against Gli3 and Hes-1 reveals strong Gli3 expression in the Hes-1 positive cells (Black arrow) and lack of immunodetectable Gli3 in the most basal levels (white arrow). I) Proposed model with Gli3 expression in proliferative Hes-1+ cells and in daughter cells that exit the proliferative program and enter the Ascl-1+ controlled pro-neurogenic program in the basal regions of the developing VNO. Scale bars A,C,E,G: 100 μm, B,D,F,H: 50 μm

### Gli3 loss-of-function affects VSNs neurogenesis but retains VSNs’ specification and differentiation

The expression pattern of Gli3 in apical region of the VNO resembles its expression in the ventricular zone of the developing brain cortex (Hasenpusch-Theil et al., 2018). Gli3 immunoreactivity in the apical HES1+ proliferative region of the VNO prompted us to analyze if Gli3 repressor activity contributes to initiating the differentiation of neurogenic progenitors (Wang et al., 2011; Hasenpusch-Theil et al., 2018). We speculated that Gli3 may play a similar role as in the cortex to stimulate the onset of neurogenic progenitors responsible for the genesis of GnRH-1ns, terminal nerve cells, and VSNs. VSNs can differentiate into either apical or basal VSNs (Lin et al., 2018b). To identify and quantify the cell bodies of the embryonic VSNs, we performed immunolabeling against the transcription factors Meis2 and Tfap2e (AP-2ε) on Gli3^Xt/Xt^ mutants and controls (Enomoto et al., 2011; Lin et al., 2018a) (Fig.3A,B). At E15, we detected both apical and basal VSNs in the developing VNOs of controls (Fig.3A). However, we found a dramatic reduction (~70%) in the number of VSNs in Gli3^Xt/Xt^ null mutants. (Fig.3C). Heterozygous Gli3^WT/Xt^ displayed an intermediate phenotype between controls and Gli3^Xt/Xt^ mutants. These data suggest that lacking functional Gli3 compromises VSNs neurogenesis, while permitting the differentiation of the two main types of VSNs (Fig.3A-C).

**Figure 3.**
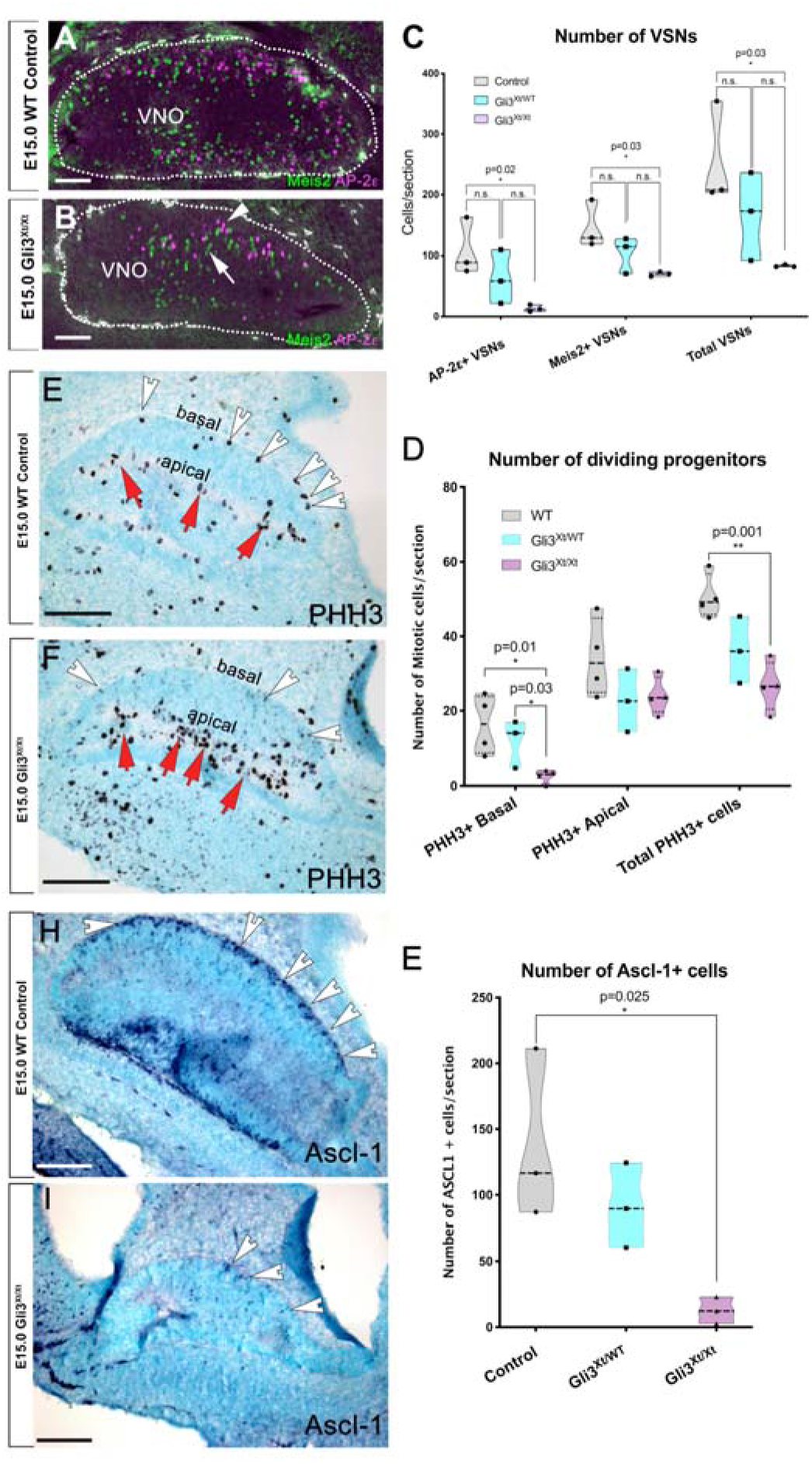
Gli3 loss-of-function impairs the formation of Ascl-1+ neuronal progenitor cells and VSN neurogenesis but not VSN terminal differentiation. A,B) Immunostaining against the transcription factors Meis2 and AP-2ε highlights maturing apical and basal vomeronasal sensory neurons in controls (A) and Gli3^Xt/Xt^ mutants (B). C) Quantifications of the average number of AP-2ε differentiated VSNs and Meis2 differentiated apical VSNs reveal a significant reduction in the number of differentiated VSNs in Gli3^Xt/Xt^ mutants. E,F) PHH3 immunostaining and quantification (D) of VNO from WT and Gli3^Xt/Xt^ reveals a reduction in the total number of dividing cells with a significant reduction of apical progenitors (white arrow), but not apical progenitors. H-I) Immunostaining against Ascl-1 in controls and Gli3^Xt/Xt^ mutants reveals (E) a significant reduction in the number of Ascl-1 positive cells in the basal domains of the developing VNO. Scale bars all are 100 μm.

Quantification of PHH3+ mitotic cells at E15 highlighted a significant decrease in the total number of mitotic progenitors in Gli3^Xt/Xt^ mutants (Fig.3E,F,D). However, we found a significant reduction in the number of basal progenitors in Gli3^Xt/Xt^ compared to WT animals and found no significant difference in apical progenitor cells (Fig.3D). Heterozygous Gli3^WT/Xt^ embryos displayed a non-significant intermediate phenotype compared to WT or Gli3^Xt/Xt^. Since we found a significant reduction in proliferation in the basal portions of the VNO, we examined if Gli3 loss-of-function compromised the onset of Ascl-1 positive basal neurogenic progenitors. Quantifying Ascl-1 positive cells confirmed a dramatic (~90%) reduction of Ascl-1+ neurogenic progenitors in the VNO (Fig.3H,I,E). These data suggest that Gli3 loss-of-function in the VNO impairs the onset of neurogenic progenitors similar to the developing cortex (Wang et al., 2011; Hasenpusch-Theil et al., 2018). We conclude that the loss of Gli3 mediated gene repression negatively affects the onset of neurogenic progenitors in the VNO.

### Gli3 loss-of-function disrupts GnRH-1 neuronal migration

The observed defects in VSN development prompted us to investigate the impact of Gli3 loss-of-function on GnRH-1 neurogenesis and migration. In control conditions, GnRH-1ns migrate along the axons of the terminal nerve that invade the brain ventral to the olfactory bulb, cross the developing ventral telencephalon/sub-pallium, then project to the preoptic area (Cariboni et al., 2011; Hanchate et al., 2012; Giacobini and Prevot, 2013; Taroc et al., 2017) (Fig.4A,C,D,G). However, Gli3^Xt/Xt^ mutants show large clusters of cells in the nasal area (Fig.4B,E), as most GnRH-1ns were unable to reach the brain. Some GnRH-1ns in these mutants migrated toward the developing brain but then formed cell clumps along the migratory track (Fig.4F). Sparse GnRH-1ns did reach the putative forebrain junction without invading the brain (Fig.4B,H).

**Figure 4.**
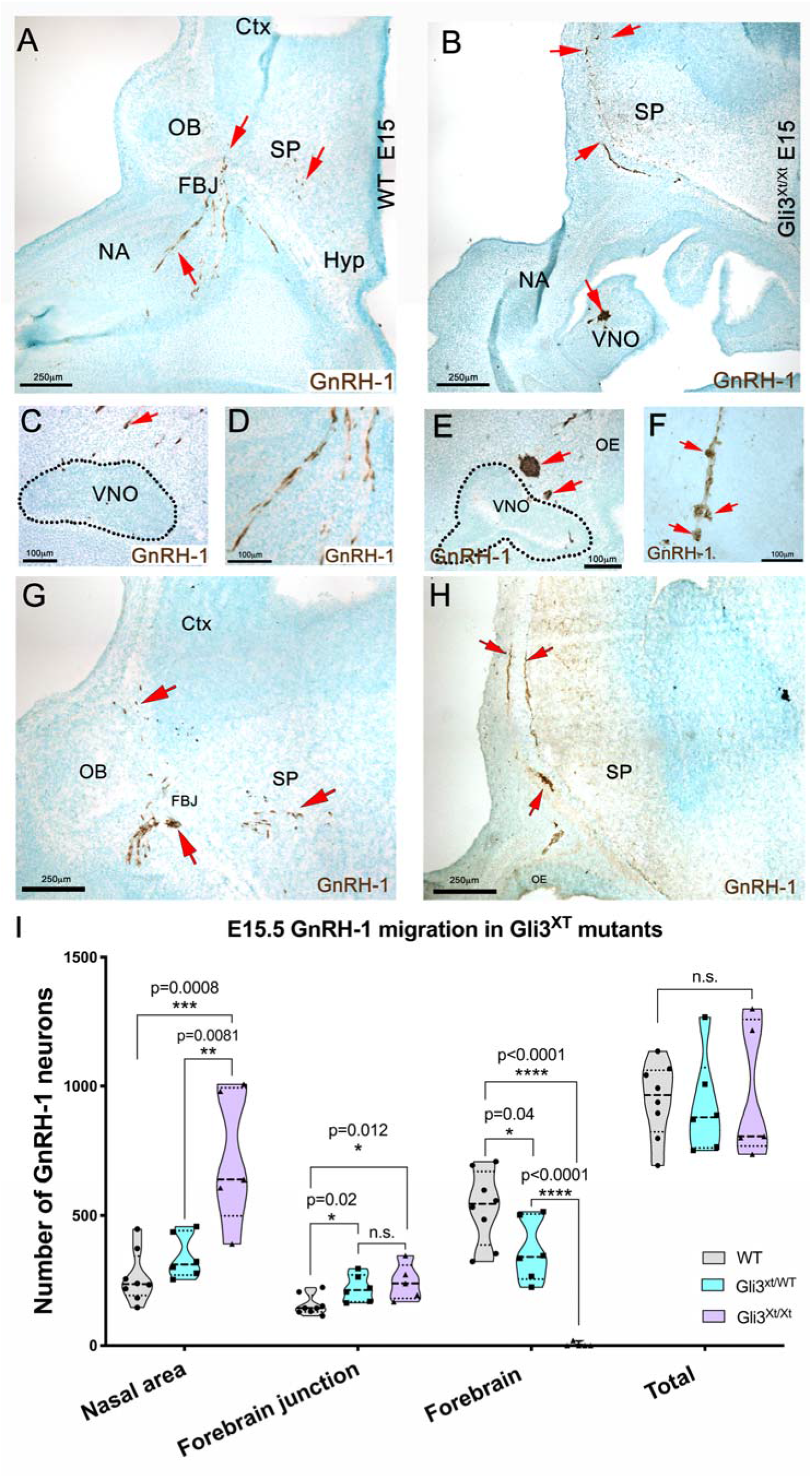
Gli3^Xt/Xt^ mutants display severely impaired GnRH-1 neuronal migration. Representative images of immunostaining against GnRH-1 on WT (A-D) and Gli3^Xt/Xt^ (E-H) at E15.5 (A) in WT controls. GnRH-1ns migrate as a continuum from the vomeronasal organ (VNO) to the brain. GnRH-1ns from the nasal area (NA), cross the forebrain junction (FBJ), invade the brain ventral to the OB, migrate through the subpallium (SP) to eventually reach the hypothalamic area (Hyp). B,C) GnRH-1ns migrating in the nasal area and out of the VNO. D) Detail showing GnRH-1ns invading the subpallium (SP) ventral to the OB in the region corresponding to the putative accumbens (Ac), some GnRH-1ns migrate to the cortex (Ctx) and around the olfactory bulbs (OBs). E-H) In Gli3^Xt/Xt^, a large portion of GnRH-1ns neurons form clusters of cells proximal to the VNO while other migrate around the brain, occasional GnRH-1ns were found accessing the brain dorsal portion of the cortex (Ctx). F,G) GnRH-1 immunostaining (brown) show less organized GnRH-1ns forming clumps while migrating towards the brain (compare to B). G) Most of the GnRH- 1ns form large clusters proximal to the VNO (arrows). H) Detail showing GnRH-1ns were unable to invade the brain migrating along the meninges. I) Quantification of the distribution of GnRH-1ns in WT, Gli3^Xt/WT^ and Gli3^Xt/Xt^. In Gli3^Xt/Xt^ the majority of the GnRH-1ns remains in the nasal area. Both Gli3^Xt/WT^ and Gli3^Xt/Xt^ have a significant smaller number of GnRH-1ns in the brain compared to controls. Values ± SE, dots indicate numbers of embryos/genotype, unpaired t-test, significant values p<0.05.

We then quantified the distribution of GnRH-1ns between the nasal area forebrain junction and brain on serial parasagittal sections after immuno-staining against GnRH-1 (Fig.4I). These data revealed a lack of GnRH-1 immunoreactive cells in the brain of Gli3 mutants with a massive accumulation of GnRH-1 cells in the nasal area. We observed a small, but significant, reduction in the number of GnRH-1 positive cells in the brain of heterozygous controls at E15 with a comparable increase in GnRH-1 cells in the nasal area. Notably, quantifying WT, Gli3^Xt/WT^, and Gli3^Xt/Xt^ mutants at E15 revealed that the total number of GnRH-1ns was comparable across genotypes. These data indicate a distinct lineage for VSNs and GnRH-1ns. We propose that Gli3 loss-of-function compromises the formation of Ascl1-1+ vomeronasal progenitors (Fig.3), but not GnRH-1 neurogenesis or differentiation.

### Gli3 loss-of-function impairs OECs development in the nasal mucosa

We investigated if the defective migration of GnRH-1ns in Gli3^Xt/Xt^ results from defective vomeronasal development and/or aberrant terminal nerve projections to the brain. So, we performed a series of immunolabeling experiments to discriminate between cell types. Immunolabeling with anti-peripherin provided a shared marker to highlight olfactory, vomeronasal, and terminal nerve neurons (Fig.5A-D,M,N)(Taroc et al., 2017). To follow the projections of VSNs and the terminal nerve (See cartoon in Fig.5; Fig.5A-D), we performed immunolabeling against the guidance receptors, Neuropilin-2 (Nrp2), Roundabout-2 (Robo2) and Neuropilin-1 (Nrp1). Robo2 highlights projections of the basal VSNs (Fig.5E,F,I,J), Nrp2 highlights apical VSNs (Fig.5G-J) (Prince et al., 2009), while Nrp1 (Fig.5K2,L2) (Taroc et al., 2017) labels projections of putative terminal nerve neurons (Fig.5K1-L2). In control animals, we found TN fibers (Nrp1+) invaded the brain, together with GnRH-1ns (Fig.5A,C,E,G,I). However, Gli3^Xt/Xt^ mutants showed stalled olfactory and vomeronasal axons (positive for Robo2 and/or Nrp2, Fig.5B,D,F,H,J) (Balmer and LaMantia, 2004) in front of the forebrain, which formed a FCM. However, we did observe that sparse putative terminal nerve axons and GnRH-1ns (Fig.5B,L1,L2) did reach the putative forebrain junction and project around the dorsal portions of the brain. However, the majority of GnRH-1ns (Fig.4E) and terminal nerve fibers (Peripherin+; Nrp1+, Fig.5 O1-O4; negative for Nrp2 and Robo2, Fig.5N-N4) formed large tangles in the nasal area.

**Figure 5.**
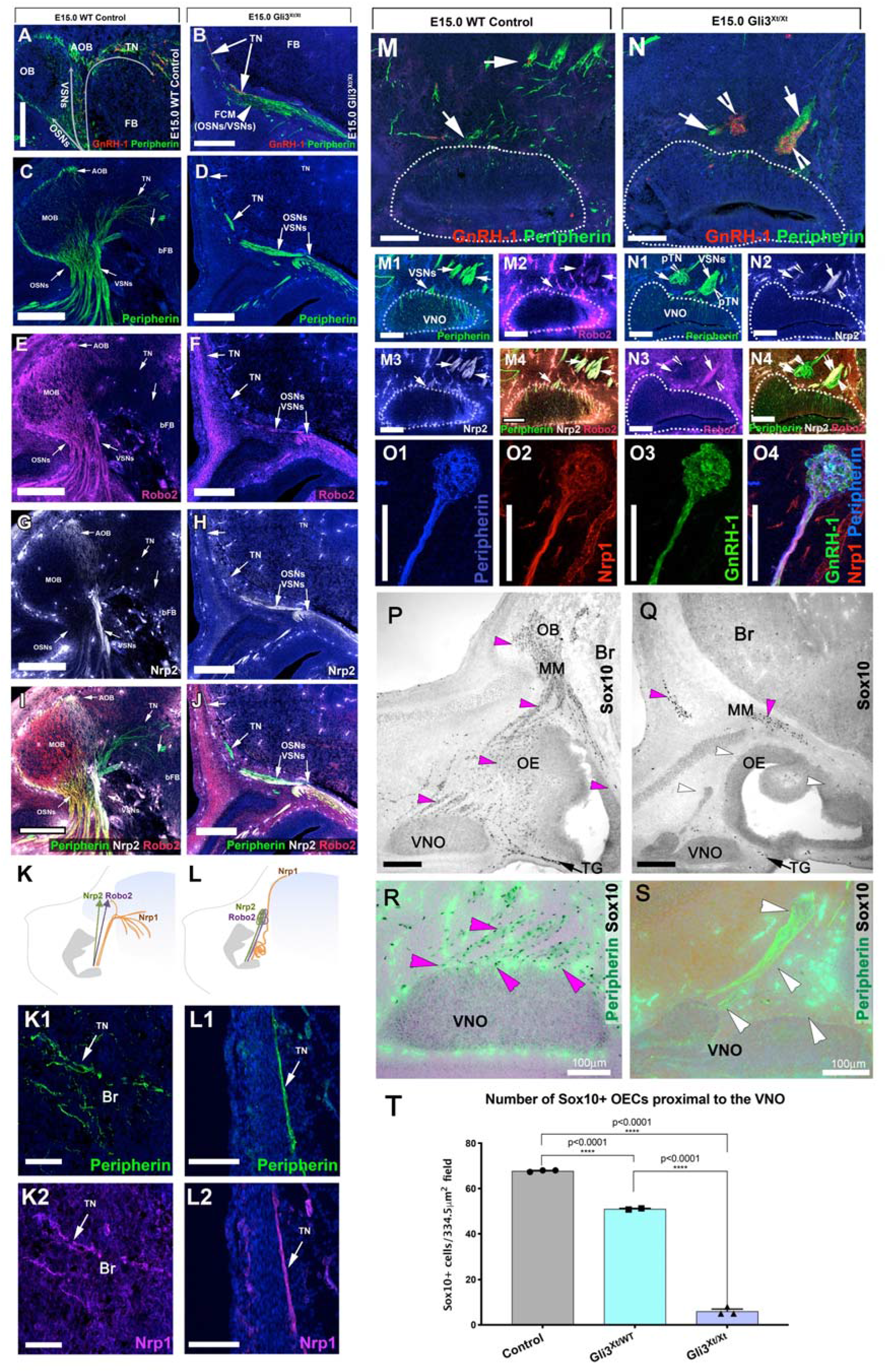
Misrouted vomeronasal and terminal nerve axons in Gli3^Xt/Xt^ are associated with lack of ensheathing cells in the nasal mucosa. A,C) Immunostainings against Peripherin and GnRH-1. B,D) Peripherin and GnRH-1 immunofluorescence, in Gli3^Xt/Xt^ mutants the OSNs and VSNs form a fibrocellular cellular mass (FCM). The GnRH-1ns (D) and the TN do not remain in the FCM but migrate around the FB. C-I) WT E15.5, immunostaining against Peripherin, Robo2, Nrp2; the TN fibers are negative for Robo2 expression and negative for Nrp2 (see merge in I). D-J) Gli3^Xt/Xt^ E15.5 immunostaining against Peripherin, Robo2, and Nrp2 and merge. K,L) Schematic of the phenotypes and fiber immunoreactivity in controls and Gli3^Xt/Xt^ mutants. K1,K2) Immunofluorescence showing Neuropilin-1 (Nrp1) immunoreactivity in TN fibers invading the brain in WT embryos. L1,L2) Peripherin and Neruopilin1 positive fibers of the putative TN project around the brain. M-M4) In control animals the GnRH-1ns migrate along Peripherin positive bundles (M1), also immune reactive for Robo2 (M2), Nrp2 (M3), Merge in M4. In Gli3^Xt/Xt^, GnRH-1ns form large clumps of cells suggesting disorganized Peripherin positive fibers. Some tangled fibers of the putative terminal nerve (N1) are negative for Nrp2 (N2) and Robo2 (N3). Merge in N4. O1-O4) In Gli3^Xt/Xt^, tangles of putative TN fibers are positive for peripherin (O1) and Nrp1 (O2) and are associated with clusters of GnRH-1ns (O3). Merge in O4. P-S) GLI3^Xt/Xt^ lack of OECs associated with the vomeronasal nerve. P-S) Combined Peripherin IF and Sox10 IHC. P) In controls, Sox10 immunoreactive OECs (magenta arrowheads) were found around the VNO, proximal to the basal lamina of the OE, along the Peripherin positive vomeronasal and olfactory bundles and as part of the migratory mass (MM) in front of the brain. Sox10+ Schwann cells were found associated along nasal trigeminal projections (TG). Q) In Gli3^Xt/Xt^ mutants, Sox10 immunoreactive OECs were neither found proximal to the basal portions of the VNO, olfactory epithelium nor along the vomeronasal and olfactory projections (white arrowheads). Notably Sox10 positive cells were found in the migratory mass (MM). As in controls Sox10+ Schwann cells were found associated along nasal trigeminal projections (TG). R,S) Magnifications of the VNO. R) In control animals Sox10 positive OECs were found proximal to the basal lamina of the VNO, along the vomeronasal bundles. S) In Gli3^Xt/Xt^ Sox10+ OECs were rarely found in proximity of the basal lamina of the VNO and along the vomeronasal/terminal nerve bundles (white arrowheads). T) Quantification of Sox10+ OECs around the VNO, ****p<0.000. Scale bars are all 100 μm.

GnRH-1 neuronal development strictly depends on correct formation of OECs (Barraud et al., 2013; Pingault et al., 2013). Gli3 can act as a Sox10 modifier (Matera et al., 2008). Our data show that Gli3 loss-of-function leads to defective vomeronasal and terminal nerve trajectories, which phenocopies that described after OECs loss in Sox10 null mouse mutants (Barraud et al., 2013; Pingault et al., 2013). So, we tested the role of OECs in axonal misrouting and GnRH-1 migratory defects in Gli3^Xt/Xt^ mutants. So, we performed double staining against Peripherin and Sox10 (Forni et al., 2011b; Pingault et al., 2013; Rich et al., 2018). In control animals, Sox10 positive OECs were localized proximal to the basal lamina of the vomeronasal epithelium and correlated with the vomeronasal bundles emerging from the vomeronasal epithelium (Fig.5P,R). In the OE and olfactory fibers, we observed a similar distribution with strong Sox10 immunoreactivity close to the basal regions of the OE and along the olfactory bundles projecting to the OBs (Fig.5P). In Gli3^Xt/Xt^ mutants, we found a near complete lack of Sox10 immunoreactivity around the vomeronasal epithelium and no Sox10+ cells associated with vomeronasal axons or terminal nerve (Fig.5Q,S). We also found no Sox10 immunoreactivity near the basal lamina of the olfactory epithelium (Fig.5Q). However, in Gli3^Xt/Xt^, we found Sox10 positive ensheathing cells proximal to the brain as part of the FCM (Fig.5Q), consistent with previous reports (Balmer and LaMantia, 2004).

As we detected dramatic defects in OECs formation in the developing olfactory system, we propose that the aberrant neuronal patterning and GnRH-1 migratory impairment observed in Gli3^Xt/Xt^ mutants is secondary to the lack of OECs in the nasal mucosa (Barraud et al., 2013; Pingault et al., 2013; Rich et al., 2018).

### Delayed, but not defective, migration in Gli3^Xt/WT^ heterozygous mutants

Partial defects in GnRH-1 neuronal migration in Gli3^Pdn/Pdn^ Gli3 hypomorphs (Naruse et al., 1994) suggest that defects in Gli3 levels can impair GnRH-1 neuronal migration. As we collected evidence indicating that Gli3 controls neurogenesis of VSNs, we further analyzed the development of the GnRH-1 system in Gli3^Xt/WT^ heterozygous embryos. Gli3^Xt/WT^ are viable and fertile, and have mild craniofacial defects and an extra toe (Veistinen et al., 2012). Yet, they do not display dramatic abnormalities in the olfactory system. We analyzed GnRH-1 neuronal migration in Gli3^Xt/WT^ heterozygous animals after development. Analysis of adult WT controls and Gli3^Xt/WT^ mutants (Fig.6A-C) revealed no significant differences in the number and distribution of GnRH-1ns in the brain after development. Immunostaining against GnRH-1 and quantifying the average area of GnRH-1 immunoreactive axons also revealed comparable GnRH-1 innervation of the median eminence (Fig.6D-F). These data suggest that Gli3 haploinsufficiency mice show delayed, but not disrupted, GnRH-1 neuronal migration.

**Figure 6.**
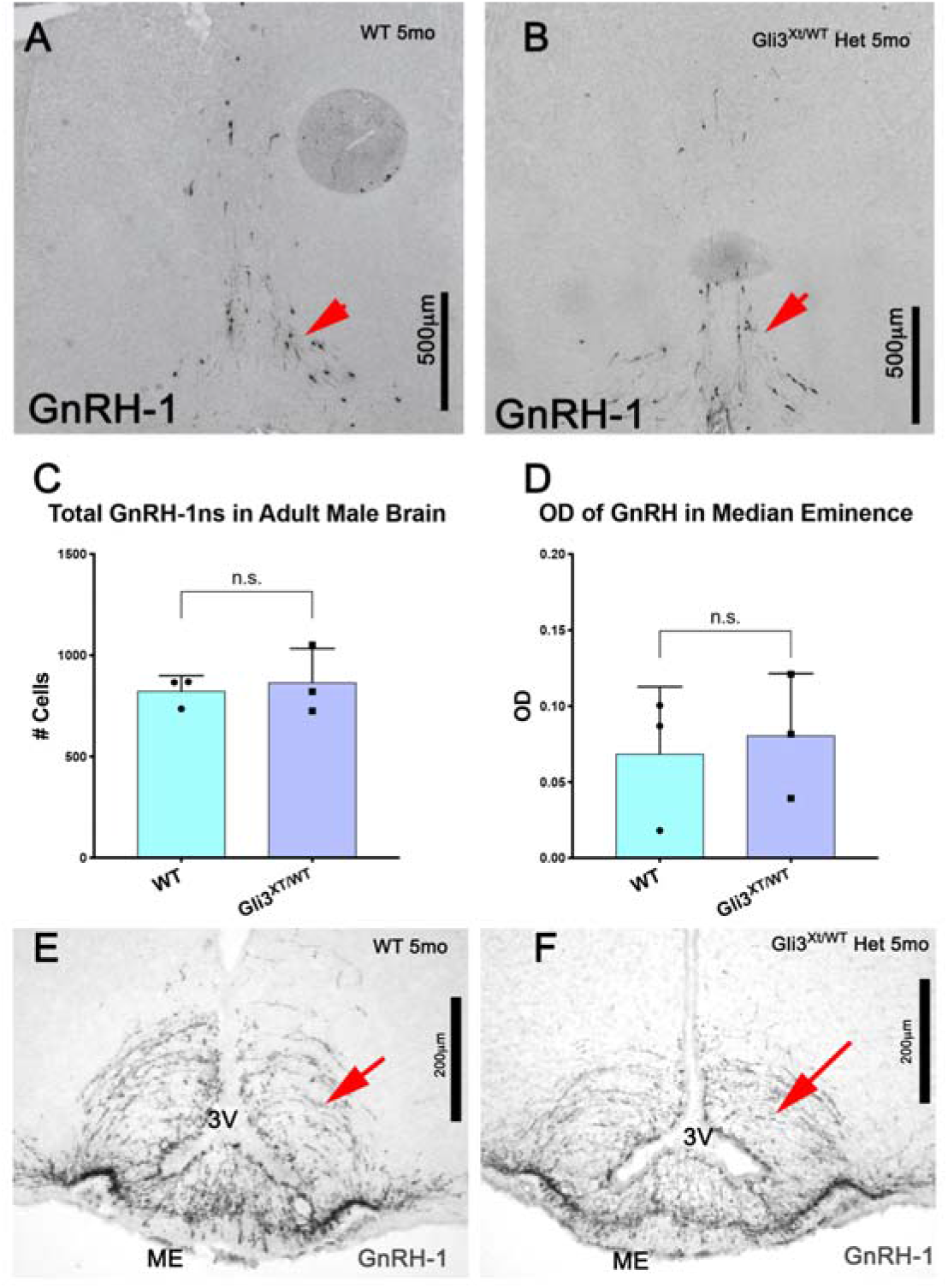
Gli3^Xt/WT^ show no difference in GnRH-1ns numbers in the brain after birth. A, B) Immunostaining against GnRH-1 on postnatal WT and Gli3^Xt/WT^ heterozygous animals shows comparable immunoreactivity in GnRH-1 cell bodies in the preoptic area (arrows). C) Quantification for GnRH-1 immunoreactive cell bodies in the preoptic area indicates no statistical differences between controls and Gli3^Xt/WT^ heterozygous mutants. D) Quantification of area occupied by GnRH-1 immunoreactive fibers in the median eminence of WT controls and Gli3^Xt/WT^ heterozygous mutants. E, F). Representative images showing GnRH-1 immunoreactivity of GnRH-1 axons in the median eminence (ME) of WT and Gli3^Xt/WT^ heterozygous mutants.

### Lack of TN and GnRH-1 invasion of the brain and altered Sema3A expression in Gli3^Xt/Xt^ mutants

In Gli3^Xt/Xt^, the few TN and GnRH-1 neurons that reach the brain fail to invade it. Since brain development can be highly dysmorphic after Gli3 loss-of-function (Aoto et al., 2002; Balmer and LaMantia, 2004; Fotaki et al., 2006; Blaess et al., 2008; Veistinen et al., 2012; Hasenpusch-Theil et al., 2018), we analyzed how such aberrant development can indirectly compromise the ability of TN and GnRH-1 neurons that do reach the brain to invade it (Fig 4H; 5K,L). In rodents, gamma-aminobutyric acid (GABA)-ergic interneurons form in the ventral telencephalon and subsequently migrate toward the cortex (Anderson et al., 1997; Lavdas et al., 1999; Wichterle et al., 1999; Wichterle et al., 2001; Markram et al., 2004). The transcription factor AP-2ε can identify the developing olfactory bulbs and vomeronasal neurons (Feng et al., 2009; Besse et al., 2011; Lin et al., 2018a) (Fig.7A-D). We used antibodies against glutamate decarboxylase 1 (brain, 67kDa) (GAD67/GAD-1) to label the ventral forebrain/subpallium. Consistent with previous reports (Yu et al., 2009; Besse et al., 2011), Gli3^Xt/Xt^ mutants exhibit a lack of rostral olfactory bulb protrusions but form ectopic olfactory bulb-like structures, positive for AP-2ε (Besse et al., 2011) in the dorsal/lateral portions of the developing telencephalon (Fig.7B,D). At E15.5, GnRH-1ns invade the brain by crossing the putative accumbens (ventral to the OB) and then steering ventrally toward the hypothalamic area.

**Figure 7.**
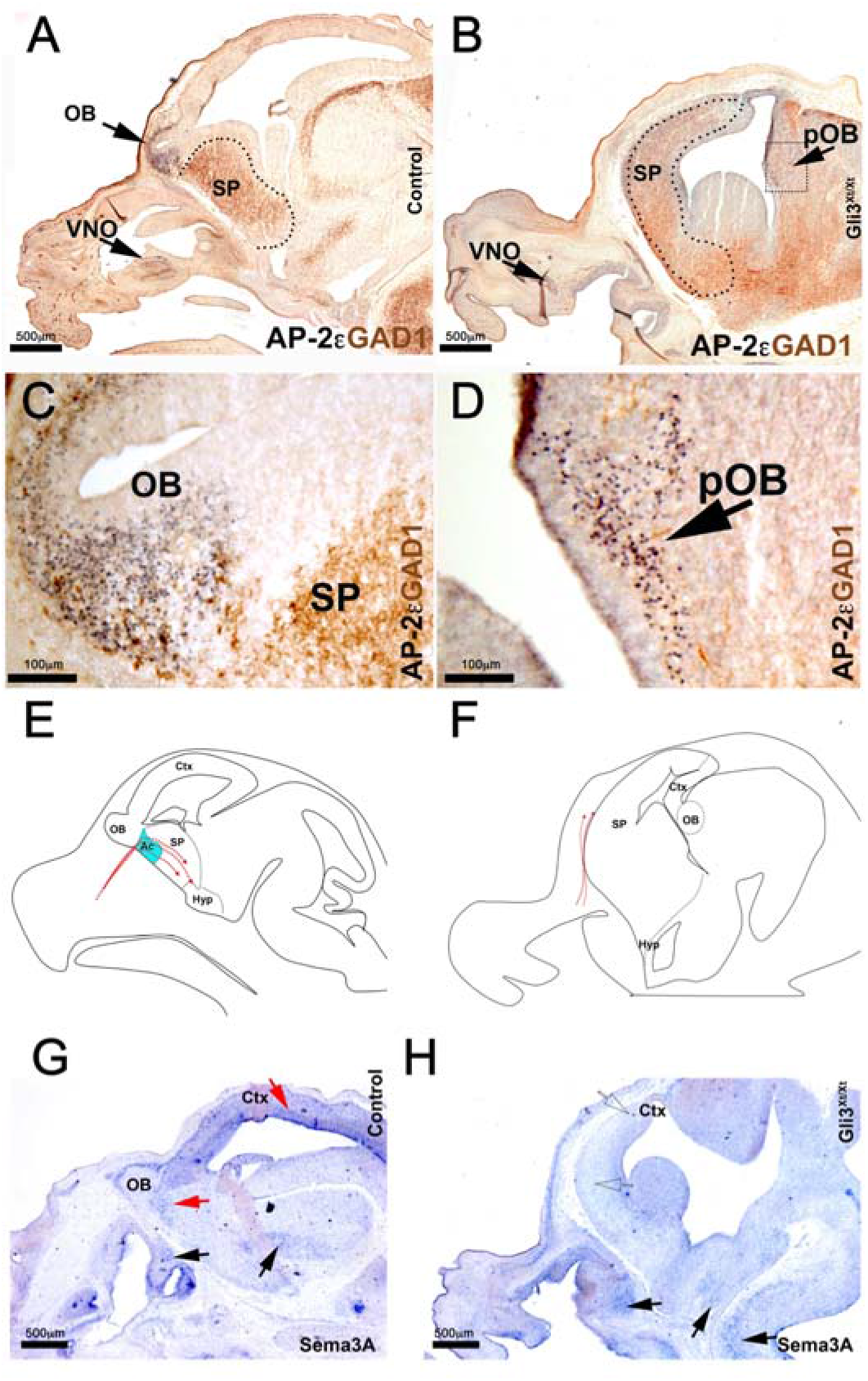
Gli3^Xt/Xt^ mutants display an expansion of expression of forebrain ventral markers, form ectopic olfactory bulbs and lack of Sema3A expression in the forebrain. A) AP-2ε/GAD1 double immunostaining. Immunostaining against AP-2ε highlights mitral cells in the OB (magnified in C). GAD1 immunostaining highlights GABAergic neurons which are mostly located in the subpallium. B) In Gli3^Xt/Xt^, AP-2ε highlight extopic mitral cells in the posterior portion of the forebrain (boxed, see magnification in (D) showing the ectopic AP-2ε+ OB in the posterior portion of the brain. GAD1 expression shows expansion of ventral forebrain maker. E-F) Illustration of GnRH-1ns migration in WT and Gli3 null animals. Area of Sema3A expression represented in light blue present in WT accumbens absent in the Gli3 KOs. G-H) In-situ hybridization against Sema3A shows detectable Sema3A expression in the nasal area, basal brain (black arrows) and hindbrain of control and Gli3 mutants. No Sema3A expression was detected in Gli3^Xt^ mutants in the forebrain and cortex (compare red arrows in I with dotted arrows in H). Ac=accumbens, Ctx=cortex, Hyp=hypothalamus, OB=olfactory bulb, pOB=putative olfactory bulb, SP=subpallium.

During embryonic development, Sema3A is expressed ventral to the OB in the putative accumbens/ “septostriatal transition area” (Fig.7E,G)(Taroc et al., 2017). Lack of Sema3A is not compatible with normal invasion of the brain by the TN and GnRH-1ns (Cariboni et al., 2011). As the development of the brain appears highly dysmorphic in Gli3 mutants (See cartoon in 7F), we analyzed the pattern of Sema3A expression compared to controls. Our analysis revealed that Gli3^Xt/Xt^ mutants showed comparable Sema3A expression to controls in nasal areas and the hindbrain. However, we found a near complete absence of Sema3A expression in the expanded subpallium and the dorsal portion of the forebrain (Fig.7B,D). These data suggest that Gli3 loss-of-function compromises Sema3A expression and patterning in the brain, which, alone, is a molecular condition sufficient to prevent migratory GnRH-1 neurons to invade the brain (Cariboni et al., 2011; Taroc et al., 2017).

### Ascl-1 loss-of-function compromises VNO and GnRH-1 neurogenesis but retains OECs formation in the nasal mucosa

The negative effects of Gli3 loss-of-function on VSNs’ but not on GnRH-1ns’ neurogenesis prompted us to test if 1) Ascl1 helps control GnRH-1 neurogenesis (Kramer and Wray, 2000; McNay et al., 2006) and 2) the lack of Sox10+ OECs in the mucosa and GnRH-1 migratory phenotype observed in Gli3 mutants directly reflects compromised vomeronasal neurogenesis (Cau et al., 1997; Cau et al., 2002). We analyzed GnRH-1, VSN, and OECs development at E13.5. At this developmental stage, the TN and VSNs form Peripherin positive fiber bundles that project toward the brain. We found no effects after loss of one Ascl1 allele on vomeronasal neurogenesis or the number and distribution of GnRH-1ns (Data not shown). However, in Ascl-1 KOs we found a (~60%) reduction in VSNs compared to controls which is similar that in Gli3^Xt/Xt^ (Fig.8 I,J,O). As for the reduced neurogenesis, in Ascl-1 null animals we observed a similar (60% ; SE +/−1%; p=0.003) reduction in the number of OECS (Fig.8 K-N,O). However, SOX10+ OECs were found to be distributed with similar frequency to the controls along the vomeronasal projections (WT 0.080 cell/μm n=3; Ascl1 KO= 0.082 cell/μm, n=3; p=0.89). In Ascl-1 KOs, we found sparse GnRH-1 immunoreactive neurons emerging from the developing VNO and migrating toward the brain (Fig.8B,D,F,H). Cell quantifications revealed a significant (~86%) decrease in the total number of GnRH-1 immunoreactive neurons in Ascl-1 KOs (Fig.8P). As we found no defect in Ascl1+/− but in Ascl1 KOs, our data indicate that Ascl1 has no dose dependent effect on vomeronasal and GnRH-1 neurogenesis (Data not shown).

**Figure 8.**
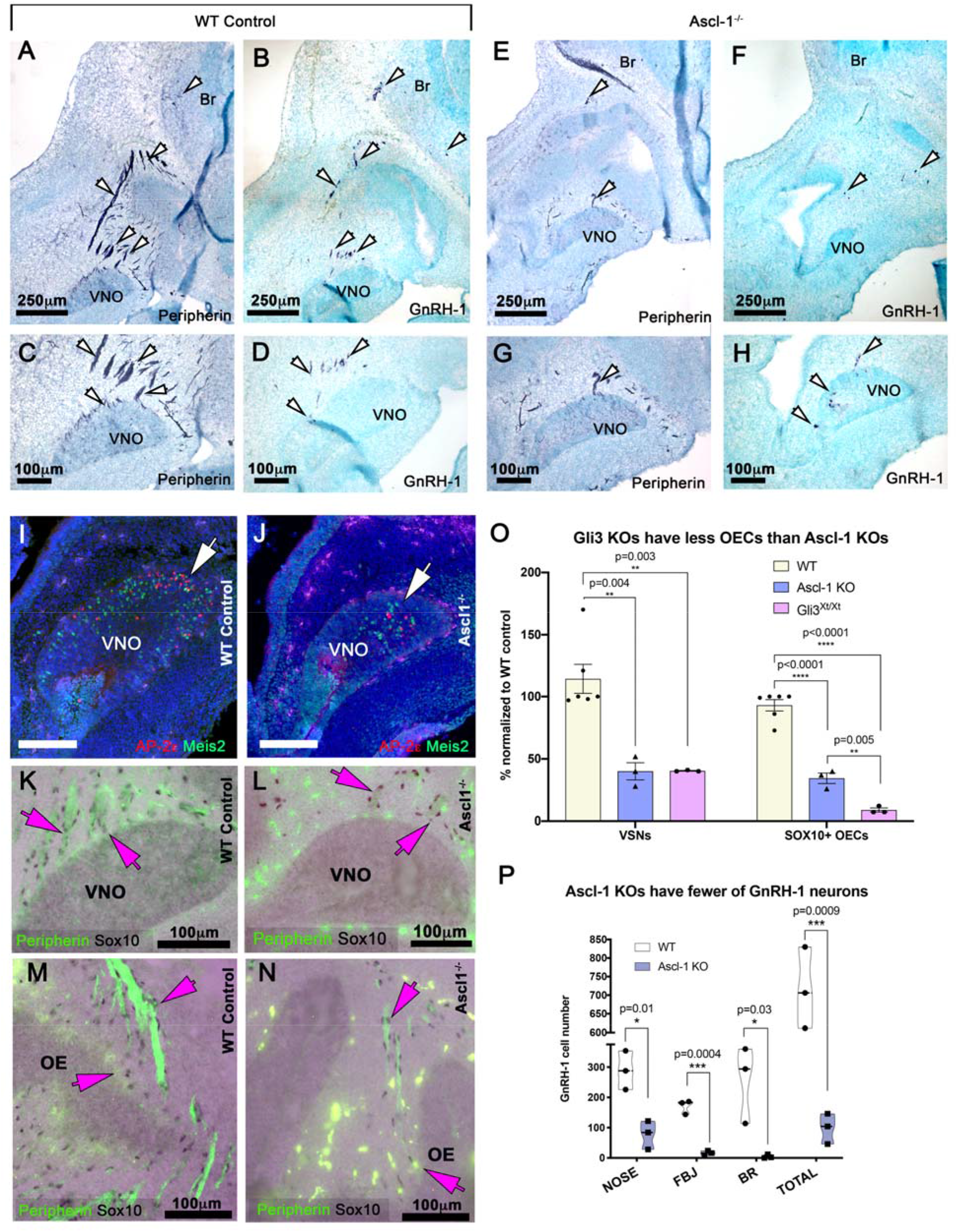
Ascl-1 KO have a comparable reduction in the number of vomeronasal and GnRH-1 neurons. A-C) Immunostaining against Peripherin highlights neuronal projections (white arrows) emerging from the VNO of WT animals projecting towards the brain. B, D) In controls, immunostaining against GnRH-1 reveals migratory GnRH-1ns distributed between the VNO and the brain. E, G) In Ascl-1 null mutants, only sparse peripherin positive projections emerge from the OE and VNO (white arrows). F-H) In Ascl-1 KOs, GnRH-1ns emerge from the VNO migrating in the nasal area (white arrows). I, J) AP-2ε and Meis2 immunostaining highlights the reduced number of differentiated VSNs in Ascl-1 KOs (white arrow). K-N) Reduced olfactory and vomeronasal neurons in Ascl-1 KOs does not prevent ensheathing cell formation in the nasal mucosa. K-N) Combined IF anti Peripherin and Sox10 IHC. K, M) In control animals, many Sox10 immunoreactive OECs (magenta arrows) are found around the VNO, along the Peripherin positive vomeronasal bundles, (M) proximal to the developing OE and along the olfactory bundles. L) In Ascl-1^−/−^ mutants, Sox10 immunoreactive OECs were found associated with the sparse vomeronasal projections (magenta arrows) (N) proximal to the developing OE and along the olfactory bundles (magenta arrows). P) Quantification of VSNs, OECs, show similar defects in vomeronasal neurogenesis between Ascl-1^−/−^ and Gli3^Xt/Xt^ compared to WT controls however, more severe loss of OECs was found in Gli3^Xt/Xt^ compared to Ascl-1^−/−^. O) Quantification of GnRH-1ns in controls and Ascl-1 KOs shows dramatic reduction in the number of GnRH-1ns in Ascl-1 KOs. Forebrain junction (FBJ). Holm-Sidak’s multiple comparisons test. White scale bars are 100 μm.

As we observed a severe reduction in the number of GnRH-1ns in Ascl-1 null, but not in Gli3^Xt/Xt^, mutants, our data suggest that the mechanisms controlling Ascl-1 expression in GnRH-1 neuronal progenitors are independent of Gli3 control. Further these observations suggest that the near complete absence of Sox10+ OECs in the nasal mucosa of Gli3^Xt/Xt^ mutants is not a direct reflection of defective vomeronasal neurogenesis/maturation (Fig.8O) (Miller et al., 2018; Rich et al., 2018), but likely secondary to neural crest defects (Matera et al., 2008). In support of this conjecture, we observed that Gli3^Xt/WT^/Ascl1^+/−^ double mutants were undistinguishable from Gli3^Xt/WT^ single mutants (Data not shown).

### A*GLI3* loss-of-function mutation in a KS and polydactyl patient

Since KS features reproductive failure and various defects in GnRH-1 neuronal migration, we first examined the genetic constraint for *GLI3* gene in Genome Aggregation Database (gnomAD) v.2.1 (Karczewski et al., 2019). This population genetics data showed that *GLI3* human genetic variants were highly constrained for loss-of-function variants with a *pLI* of 1.0 and an observed/expected (*oe*) metric of 0.09 (upper bound *oe* confidence interval of 0.20). The suggested threshold for mendelian case analysis is an upper bound *oe* confidence interval of <0.35. These parameters suggest an extreme intolerance to loss-of-function variants reflecting the strength of selection acting on heterozygotes. So, we queried WES data from 632 KS probands for *GLI3* loss-of-function variants. We identified two qualifying *GLI3* loss-of-function variants from WES data. Sanger sequencing confirmed one novel heterozygous *GLI3* loss-of-function (frameshifting indel) variant (c.1161del; p.P388Qfs*13; minor allele frequency in gnomAD: 0) in a KS male subject. Since parental DNA samples were not available, the precise mode of inheritance of the variant could not be evaluated. Additional KS/nIHH gene analysis showed that this KS proband also harbored a heterozygous *GNRHR* missense mutation (p.Q106R) (Fig.9). Since gnomAD variants are not Sanger-confirmed, we performed a gene-based burden test between the KS cohort and gnomAD for all qualifying *GLI3* loss-of-function variants that showed enrichment in the KS cohort (p=0.002). We did not observe this enrichment when correcting for Sanger-confirmed variants only in the KS cohort but retaining all unconfirmed gnomAD variants (p=0.06).

**Fig. 9.**
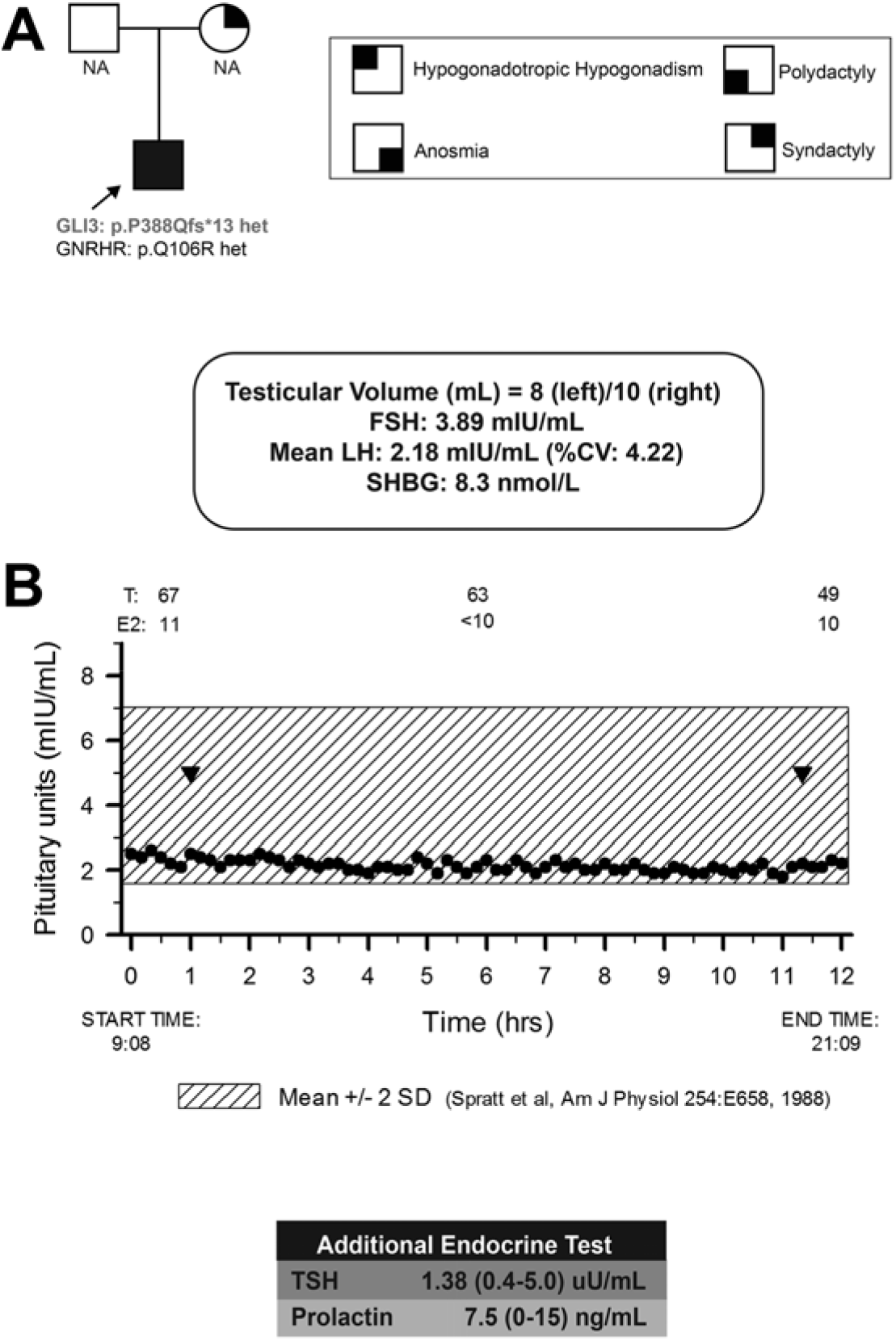
Panel A. Pedigree of Kallmann Syndrome proband (shown with arrow) with reproductive and non-reproductive phenotypes. Panel B. KS proband underwent neuroendocrine profiling (Q 10 min X 12hrs) to chart GnRH-induced LH secretion showing low amplitude, low frequency pulses (inverted triangles indicate LH pulses). Shaded region represents the normal reference range. NA, Not available; T, total testosterone; LH, luteinizing hormone; FSH, Follicle stimulating hormone, SHBG, sex-hormone binding globulin, TSH, thyroid stimulating hormone.

The patient harboring the *GLI3* loss-of-function variant is a 38-year-old male, who presented to medical attention at age 18 years with anosmia and failure to enter puberty. He was undervirilized with a bilateral low testicular volume (Fig.9). His biochemical investigations showed frank hypogonadotropic hypogonadism (serum testosterone: 67 ng/dL; LH: 2.18 mIU/mL and FSH: 3.89 mIU/mL) with neuroendocrine studies revealing low-amplitude low frequency LH pulse profile consistent with hypogonadotropism (Fig.9). The remaining pituitary hormone profile was normal and UPSIT showed complete anosmia. These results confirmed a clinical diagnosis of Kallmann Syndrome. Notably, in line with the well-established role of GLI3 in limb development, he reported in his history an extra toe on his right foot, for which he underwent surgery as a child, and bilateral webbed toes (2^nd^ and 3^rd^ toes). Although the precise inheritance mode of the variant was not determined due to lack of parental samples, his mother also exhibited syndactyly affecting her feet, suggesting a possible autosomal dominant mode of inheritance with variable penetrance for the reproductive phenotype (Fig.9).

To test the biological effects of the identified GLI3 variant, we performed a luciferase assay (Fig.10) to compare the activity of Wild type (WT) GLI3 and mutated GLI3 using a reporter plasmid containing wildtype GLI binding site linked to firefly luciferase (WTGliBS-Luc). A reporter carrying a mutated/nonfunctional (MutGLiBS-Luc) Gli3 binding site linked to firefly luciferase (Sasaki et al., 1997; Saitsu et al., 2005) served as negative control. In contrast with a prior report (Krauss et al., 2008), luciferase assay on HeLa cells (ATCC CCL-2) revealed that Gli3 acts a transcriptional repressor. This analysis demonstrated the deleterious loss-of-function nature of our novel *GLI3* frameshift mutation (p.P388Qfs*13), which affected the repressor ability of Gli3, consistent with a GLI3 loss-of-function as a putative mechanism for GLI3-related KS.

**Figure 10.**
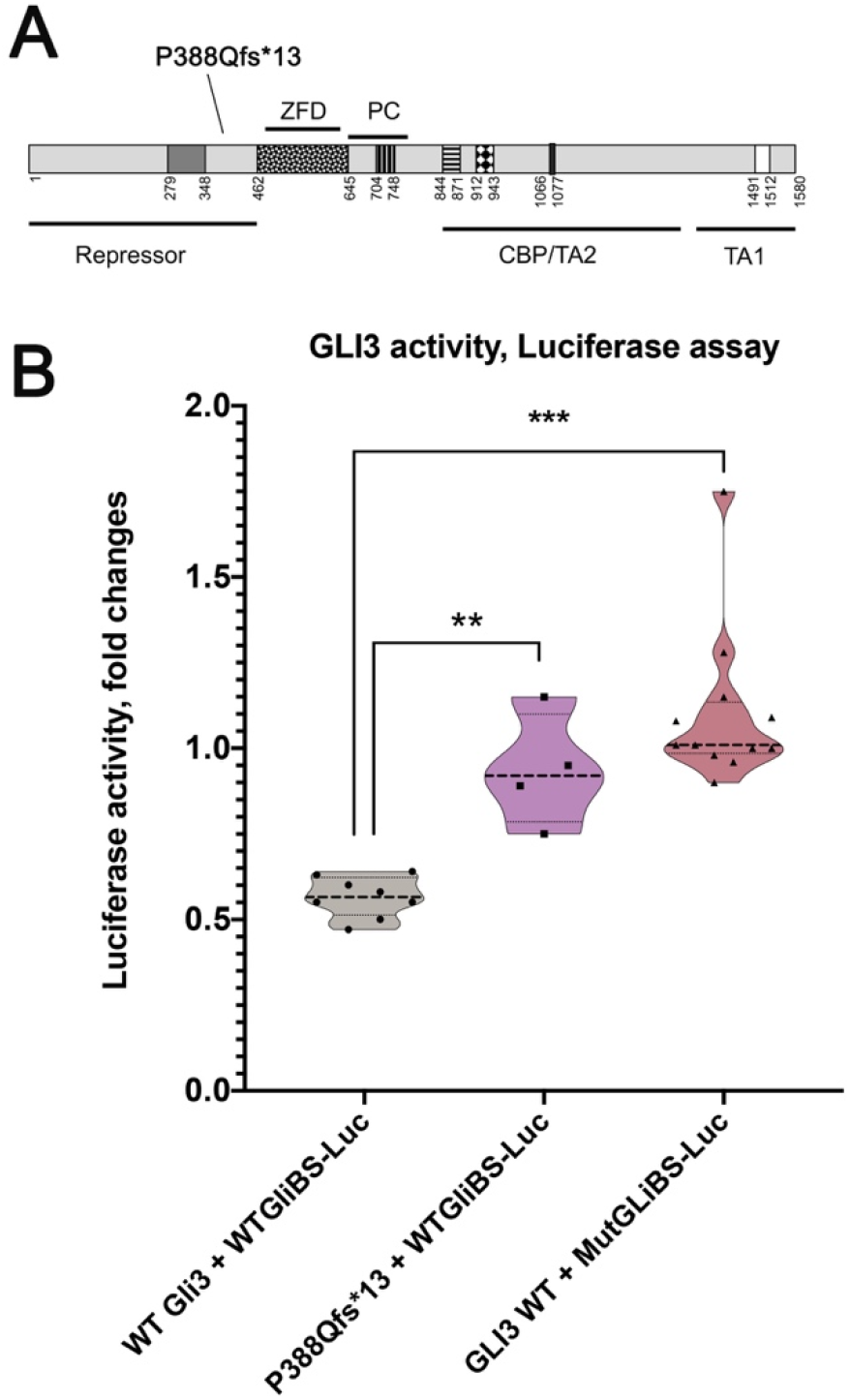
Functional validation of GLI3 loss-of-function. Cartoon illustrating *GLI3* protein, the mutation P388Qfs*13 affects the repressor domain of the protein. Zinc finger domain (ZFD), protein cleavage (PC), CBP-binding domain, transactivation domain 1 and 2 (TA1, TA2). B) Luciferase activity assay, the repressor activity of Gli3WT on the reporter (WTGLiBS-Luc) is lost in the truncated GLI3. No repression was found for WT Gli3 on reporter carrying non-functional Gli binding site MutGLiBS-Luc. **p=0.002, ***p<0.0001, Holm-Sidak’s multiple comparisons test.

## Discussion

Here, we found that Gli3 loss-of-function induces: 1) loss of Ascl-1 neuronal progenitors in the VNO and dramatically reduces the number of vomeronasal sensory neurons (VSNs), 2) normal neurogenesis for GnRH-1 neurons, 3) nearly complete absence of GnRH-1 neuronal migration, associated to aberrant formation of OECs in the nasal mucosa, 4) loss of forebrain expression of Semaphorin 3A (Sema3A), a key axonal guidance molecule that controls GnRH-1 invasion of the brain (Cariboni et al., 2011; Hanchate et al., 2012; Young et al., 2012). To elucidate the underlying cellular substrates of the observed phenotypes, we analyzed Ascl-1^−/−^ mutants. Even though we found reduced vomeronasal neurogenesis similar to Gli3 mutants, Ascl-1^−/−^ mutants showed a severely reduced number of GnRH-1ns’ number and Sox10+ OECs (OECs) that formed along axonal bundles similar to controls (Fig.8). Based on the observed phenotypes between Gli3 and Ascl1 mutants, we conclude that Ascl-1 is crucial for GnRH-1 neurogenesis independent of Gli3 control. Ascl-1^−/−^ mice showed reduced neurogenesis in the VNO alone does not prevent the maturation of Sox10 positive OECs in the nasal mucosa and the migration of GnRH-1ns in Gli3 mutants. These data suggest a direct role for Gli3 in controlling OECs development. These murine observations highlight a critical role for Gli3 distinctly in OEC development and GnRH neuronal migration, two features that underlie the pathogenesis of KS. In keeping with this notion, we also identified and functionally validated a novel GLI3 loss-of-function mutation in a KS patient who also exhibited polydactyly, a well-recognized phenotypic feature of GLI3-related human diseases.

Analyzing Gli3 expression in the developing olfactory pit, we observed that, Gli3 is expressed in proliferative Hes-1 positive cells as shown in the cortex (Hasenpusch-Theil et al., 2018). Hes-1 represses Ascl-1 expression in undifferentiated/proliferative neuroepithelial cells (Ohtsuka et al., 1999; Cau et al., 2000; Ohtsuka et al., 2001). Analyzing developing VSNs in Gli3 null mutants revealed a dramatic reduction for both apical and basal VSNs secondary to a near complete loss (~91%) of Ascl-1 positive progenitors in the VNO. Surprisingly, the number of GnRH-1ns did not appear different from WT controls. These results indicate that loss of Gli3 mediated gene repression negatively regulates the onset of vomeronasal neurogenic progenitors, but not GnRH-1 neurogenesis.

The cyclin-dependent kinase gene *Cdk6* was recently identified as a direct target of Gli3 mediated repression (Hasenpusch-Theil et al., 2018). Loss of Gli3 expression in cortical neuroepithelial cells elevates Cdk6 levels, which shortens the cell cycle and reduces neurogenesis (Pilaz et al., 2009; Hasenpusch-Theil et al., 2018). Since we detected Gli3 in Hes-1 proliferative cells, we hypothesize that Gli3 controls the onset of neurogenesis through a similar mechanism as in the cortex (Hasenpusch-Theil et al., 2018). Following Shh expression, Gli3 transitions from being a transcriptional repressor to a transcriptional activator. We performed Gli1 immunostaining on the developing VNO and did not find Gli1 expression (Data not shown). This finding suggests that Gli3 acts as a transcriptional repressor in the developing VNO. Notably, Gli3 heterozygous mutant mice showed a small delay in GnRH-1 migration during development without differences in cell number or distribution after birth and were fertile. These data indicate that heterozygous Gli3 loss-of-function mutations, alone, cannot produce a GnRH-deficient state in mice, which is consistent with results from Gli3 hypomorphic mutant mice (Naruse et al., 1994).

OECs are a glial population that wraps bundles of olfactory and vomeronasal bundles (Rich et al., 2018). OECs promote normal GnRH-1 neuronal migration from the nasal area to the brain (Barraud et al., 2013; Geller et al., 2013; Pingault et al., 2013). OECs arise from the neural crest precursors (Barraud et al., 2010; Forni et al., 2011b; Miller et al., 2016; Miller et al., 2018; Rich et al., 2018). By analyzing GnRH-1ns migration in Gli3 mutants, we found that most GnRH-1 cells were unable to migrate far from the VNO and formed large clumps of cells on tangled fibers of the putative TN. This phenotype matches that of Sox10 null mutants (Barraud et al., 2013; Pingault et al., 2013). Gli3 can act as a Sox10 modifier, which is consistent with the suggestion that Gli3 loss-of-function can compromise OECs development (Matera et al., 2008). Here, we found virtually no Sox10 positive cells in the olfactory/vomeronasal bundles of Gli3^Xt/Xt^ mutants nor proximal to the basal lamina and lamina propria of the olfactory epithelia, suggestive of Gli3 loss impairing normal OECs development in the nasal mucosa. In Gli3^Xt/Xt^ mice, we found Sox10+ Schwann cells around the trigeminal nerve (Fig.5) and Grueneberg ganglion (data not shown). We also found Sox10 positive OECs surrounding the FCM proximal to the olfactory bulbs. These intriguing data suggest developmental differences between OECs of the nasal mucosa and olfactory bulb OECs (Richter et al., 2005; Ito et al., 2006; Mayeur et al., 2013).

In Gli3 mutants we found sporadic TN fibers and GnRH-1 cells could reach the brain but not invade it. These findings revealed a similar cellular behavior to that described after Sema3A loss-of-function. Analyses of Gli3^Xt/Xt^ mutants confirmed very dysmorphic forebrain structures with expanded regions positive for the early subpallial marker, GAD1, and ectopic olfactory bulbs (Balmer and LaMantia, 2004; Fotaki et al., 2006; Blaess et al., 2008; Veistinen et al., 2012). In-situ hybridization against Sema3A showed a near complete absence of Sema3A expression in the forebrain, a condition that alone is not permissive for normal GnRH-1 migration (Cariboni et al., 2011). Taken together, we propose that altered Sema3A expression in Gli3 mutants directly reflects either the lack of functional Gli3 transcriptional control or overall aberrant brain development.

Analyzing Ascl-1 null mutants revealed a role for Ascl-1 in controlling GnRH-1 neuronal development, since we found an approximate 86% reduction in the number of GnRH-1 neurons in these mice. We also found no vomeronasal defects together with normal GnRH-1 cell numbers and migration. This finding indicates that Ascl1 is not required in a dose-dependent manner for vomeronasal and GnRH-1 neurogenesis as described for other neurons (McNay et al., 2006). The lack of overlapping phenotype between Gli3^Xt/Xt^ (normal GnRH-1 but reduced VSN cell numbers) and Ascl-1 null mutants (reduced GnRH-1 and VSNs cell numbers) indicates that GnRH-1 neurogenic onset is not regulated in a Gli3 dependent fashion.

We observed that OECs defects in Ascl1 mutants appeared less severe than in Gli3^Xt/Xt^ and OECs in the mucosa of Ascl1 mutants were positive for Sox10 (Fig. 8O). Moreover, Gli3^Xt/WT^/Ascl1^+/−^ double mutants did not have exacerbated phenotypes compared to Gli3^Xt/WT^ suggesting that the effects seen in both Ascl-1 null and Gli3^Xt/Xt^ are independent of each other and not additive (data not shown). Taken together, we conclude that the near complete absence of Sox10+ immunoreactivity in the nasal mucosa of Gli3^Xt/Xt^ mutants is not secondary to the decreased number of vomeronasal fibers (Miller et al., 2018).

We identified a novel *GLI3* frameshifting loss-of-function variant in a KS proband. Using site-directed mutagenesis and luciferase assays, we confirmed a loss of repressor ability of this mutation, which decreased the repressor ability of GLI3. This patient also displayed polydactyly and syndactyly, two hallmark clinical features of humans with *GLI3*-Greigs cephalopolysyndactyly syndrome (GCPS), which corresponds to the loss of GLI3 transcriptional repression during limb development (Johnston et al., 2010; Al-Qattan et al., 2017). Further, the identified *GLI3* loss-of-function variant (p.P388Qfs*13) affects the 5’ part of the *GLI3* gene that leads to a truncation upstream of the zinc finger domain and is well-correlated with GCPS phenotype (Johnston et al., 2010; Al-Qattan et al., 2017). Reviewing previously reported *GLI3* mutation subjects with GCPS or oral-facial-digital syndrome indicates that a small minority of these patients also display micropenis (Kroisel et al., 2001; Johnston et al., 2010), cryptorchidism (Kroisel et al., 2001; Johnston et al., 2010), or anosmia (Johnston et al., 2010). These phenotypes are consistent with an underlying KS. KS/HH may be under-recognized in these reports, as reproductive phenotypes cannot be assessed in pre-pubertal subjects harboring *GLI3* mutations. Our findings implicate heterozygous *GLI3* loss-of-function mutations as a contributing genetic factor for KS/HH, which is further supported by the associated *GLI3* missense variants in two subjects with HH (Quaynor et al., 2016).

We acknowledge that heterozygous *GLI3* mutations alone may be insufficient to cause KS/HH and may require additional genetic or non-genetic modifiers. Indeed, KS has a well-established oligogenic basis (Sykiotis et al., 2010). Here, we found that the proband with *GLI3* loss-of-function mutation also harbored an additional rare *GNRHR* p.Q106R missense mutation. Although the *GNRHR* p.Q106R variant can be deleterious in cellular assays (Leanos-Miranda et al., 2005), *GNRHR* mutations typically cause autosomal recessive hypogonadotropic hypogonadism (Gianetti et al., 2012). So, the co-occurrence of *GLI3* and *GNRHR* variants in this proband suggests a potential oligogenic (*GLI3/GNRHR*) contribution to KS.

In summary, we revealed a crucial role for Gli3 in controlling vomeronasal sensory neurons neurogenesis, formation of OECs in the nasal mucosa and GnRH-1 neuron migration. Our findings suggest a role for Gli3 in triggering the transition from a proliferative to neurogenic program in the developing VNO. Since we found a dramatic reduction in the number of Sox10 expressing OECs and aberrant Sema3A in the brain, we propose that Gli3 acts as a genetic modifier and contributes to the oligogenic nature of KS/nIHH. Future studies will delineate the gene regulatory network controlled by the Gli3 transcriptional network in the developing nasal olfactory system and identify KS/nIHH candidate genes to confirm the role of this transcription factor as a modifier in the oligogenic nature of KS/nIHH.

## Author contributions

ET, AN, JL, NP, EG, GF, conducted the experiments and performed data analysis, PF and GF performed data analysis designed the experiments, ET, RB and PF wrote the paper, GF, RB and DK shared reagents, performed data analysis and edited the paper.

## Acknowledgments

We thank Dr. Hirotomo Saitsu (Hamamatsu University School of Medicine, Hamamatsu, Japan) for sharing Gli3 luciferase reporter plasmids and Dr. Susan Wray for sharing the anti GnRH-1 (SW) antibody.

Research reported in this publication was supported by the Eunice Kennedy Shriver National Institute of Child Health and Human Development of the National Institutes of Health under the Award Numbers 1R15HD09641101 (PF), and 1R01HD097331-01 (PF) and by the National Institute of Deafness and other Communication Disorders of the National Institutes of Health under Award Number 1R01DC017149-01A1 (PF). RB is supported by the Eunice Kennedy Shriver National Institute of Child Health and Development (K23 HD077043; R01 HD096324; P50 HD HD028138). The content of this manuscript is solely the responsibility of the authors and does not necessarily represent the official views of the National Institutes of Health.

## Notes

#### Summary of Updates

This version of the manuscript has been revised to update results introducing observations on double Gli3/Ascl1 double heterozygous animals and update our conclusions

## REFERENCES

Al-Qattan MM, Shamseldin HE, Salih MA, Alkuraya FS (2017) GLI3-related polydactyly: a review. Clin Genet 92:457–466.

Anderson SA, Eisenstat DD, Shi L, Rubenstein JL (1997) Interneuron migration from basal forebrain to neocortex: dependence on Dlx genes. Science 278:474–476.

Aoto K, Nishimura T, Eto K, Motoyama J (2002) Mouse GLI3 regulates Fgf8 expression and apoptosis in the developing neural tube, face, and limb bud. Developmental biology 251:320–332.

Balasubramanian R, Crowley WF, Jr. (2011) Isolated GnRH deficiency: a disease model serving as a unique prism into the systems biology of the GnRH neuronal network. Mol Cell Endocrinol 346:4–12.

Balmer CW, LaMantia AS (2004) Loss of Gli3 and Shh function disrupts olfactory axon trajectories. The Journal of comparative neurology 472:292–307.

Barraud P, St John JA, Stolt CC, Wegner M, Baker CV (2013) Olfactory ensheathing glia are required for embryonic olfactory axon targeting and the migration of gonadotropin-releasing hormone neurons. Biol Open 2:750–759.

Barraud P, Seferiadis AA, Tyson LD, Zwart MF, Szabo-Rogers HL, Ruhrberg C, Liu KJ, Baker CV (2010) Neural crest origin of olfactory ensheathing glia. Proc Natl Acad Sci U S A 107:21040–21045.

Besse L, Neti M, Anselme I, Gerhardt C, Ruther U, Laclef C, Schneider-Maunoury S (2011) Primary cilia control telencephalic patterning and morphogenesis via Gli3 proteolytic processing. Development 138:2079–2088.

Blaess S, Stephen D, Joyner AL (2008) Gli3 coordinates three-dimensional patterning and growth of the tectum and cerebellum by integrating Shh and Fgf8 signaling. Development 135:2093–2103.

Cariboni A, Davidson K, Rakic S, Maggi R, Parnavelas JG, Ruhrberg C (2011) Defective gonadotropin-releasing hormone neuron migration in mice lacking SEMA3A signalling through NRP1 and NRP2: implications for the aetiology of hypogonadotropic hypogonadism. Hum Mol Genet 20:336–344.

Casoni F, Malone SA, Belle M, Luzzati F, Collier F, Allet C, Hrabovszky E, Rasika S, Prevot V, Chedotal A, Giacobini P (2016) Development of the neurons controlling fertility in humans: new insights from 3D imaging and transparent fetal brains. Development 143:3969–3981.

Cau E, Casarosa S, Guillemot F (2002) Mash1 and Ngn1 control distinct steps of determination and differentiation in the olfactory sensory neuron lineage. Development 129:1871–1880.

Cau E, Gradwohl G, Fode C, Guillemot F (1997) Mash1 activates a cascade of bHLH regulators in olfactory neuron progenitors. Development 124:1611–1621.

Cau E, Gradwohl G, Casarosa S, Kageyama R, Guillemot F (2000) Hes genes regulate sequential stages of neurogenesis in the olfactory epithelium. Development 127:2323–2332.

Cuschieri A, Bannister LH (1975a) The development of the olfactory mucosa in the mouse: light microscopy. J Anat 119:277–286.

Cuschieri A, Bannister LH (1975b) The development of the olfactory mucosa in the mouse: electron microscopy. J Anat 119:471–498.

Dode C, Rondard P (2013) PROK2/PROKR2 Signaling and Kallmann Syndrome. Front Endocrinol (Lausanne) 4:19.

Dode C, Teixeira L, Levilliers J, Fouveaut C, Bouchard P, Kottler ML, Lespinasse J, Lienhardt-Roussie A, Mathieu M, Moerman A, Morgan G, Murat A, Toublanc JE, Wolczynski S, Delpech M, Petit C, Young J, Hardelin JP (2006) Kallmann syndrome: mutations in the genes encoding prokineticin-2 and prokineticin receptor-2. PLoS genetics 2:e175.

Doty RL (2007) Office procedures for quantitative assessment of olfactory function. Am J Rhinol 21:460–473.

Enomoto T, Ohmoto M, Iwata T, Uno A, Saitou M, Yamaguchi T, Kominami R, Matsumoto I, Hirota J (2011) Bcl11b/Ctip2 controls the differentiation of vomeronasal sensory neurons in mice. J Neurosci 31:10159–10173.

Feng W, Simoes-de-Souza F, Finger TE, Restrepo D, Williams T (2009) Disorganized olfactory bulb lamination in mice deficient for transcription factor AP-2epsilon. Mol Cell Neurosci 42:161–171.

Forni PE, Wray S (2015) GnRH, anosmia and hypogonadotropic hypogonadism - Where are we? Front Neuroendocrinol 36C:165–177.

Forni PE, Fornaro M, Guenette S, Wray S (2011a) A role for FE65 in controlling GnRH-1 neurogenesis. J Neurosci 31:480–491.

Forni PE, Taylor-Burds C, Melvin VS, Williams T, Wray S (2011b) Neural crest and ectodermal cells intermix in the nasal placode to give rise to GnRH-1 neurons, sensory neurons, and olfactory ensheathing cells. J Neurosci 31:6915–6927.

Forni PE, Bharti K, Flannery EM, Shimogori T, Wray S (2013) The indirect role of fibroblast growth factor-8 in defining neurogenic niches of the olfactory/GnRH systems. J Neurosci 33:19620–19634.

Fotaki V, Yu T, Zaki PA, Mason JO, Price DJ (2006) Abnormal positioning of diencephalic cell types in neocortical tissue in the dorsal telencephalon of mice lacking functional Gli3. J Neurosci 26:9282–9292.

Geller S, Kolasa E, Tillet Y, Duittoz A, Vaudin P (2013) Olfactory ensheathing cells form the microenvironment of migrating GnRH-1 neurons during mouse development. Glia 61:550–566.

Giacobini P, Prevot V (2013) Semaphorins in the development, homeostasis and disease of hormone systems. Semin Cell Dev Biol 24:190–198.

Gianetti E, Hall JE, Au MG, Kaiser UB, Quinton R, Stewart JA, Metzger DL, Pitteloud N, Mericq V, Merino PM, Levitsky LL, Izatt L, Lang-Muritano M, Fujimoto VY, Dluhy RG, Chase ML, Crowley WF, Jr., Plummer L, Seminara SB (2012) When Genetic Load Does Not Correlate with Phenotypic Spectrum: Lessons from the GnRH Receptor (GNRHR). The Journal of clinical endocrinology and metabolism 97:E1798–1807.

Guo MH, Plummer L, Chan YM, Hirschhorn JN, Lippincott MF (2018) Burden Testing of Rare Variants Identified through Exome Sequencing via Publicly Available Control Data. Am J Hum Genet 103:522–534.

Hanchate NK et al. (2012) SEMA3A, a gene involved in axonal pathfinding, is mutated in patients with Kallmann syndrome. PLoS genetics 8:e1002896.

Hasenpusch-Theil K, West S, Kelman A, Kozic Z, Horrocks S, McMahon AP, Price DJ, Mason JO, Theil T (2018) Gli3 controls the onset of cortical neurogenesis by regulating the radial glial cell cycle through Cdk6 expression. Development 145.

Hu Y, Butts T, Poopalasundaram S, Graham A, Bouloux PM (2019) Extracellular matrix protein anosmin-1 modulates olfactory ensheathing cell maturation in chick olfactory bulb development. Eur J Neurosci.

Ito D, Ibanez C, Ogawa H, Franklin RJ, Jeffery ND (2006) Comparison of cell populations derived from canine olfactory bulb and olfactory mucosal cultures. Am J Vet Res 67:1050–1056.

Johnston JJ et al. (2010) Molecular analysis expands the spectrum of phenotypes associated with GLI3 mutations. Hum Mutat 31:1142–1154.

Kagoshima M, Ito T (2001) Diverse gene expression and function of semaphorins in developing lung: positive and negative regulatory roles of semaphorins in lung branching morphogenesis. Genes Cells 6:559–571.

Karczewski KJ et al. (2019) Variation across 141,456 human exomes and genomes reveals the spectrum of loss-of-function intolerance across human protein-coding genes. bioRxiv:531210.

Keino H, Masaki S, Kawarada Y, Naruse I (1994) Apoptotic degeneration in the arhinencephalic brain of the mouse mutant Pdn/Pdn. Brain Res Dev Brain Res 78:161–168.

Kim EJ, Ables JL, Dickel LK, Eisch AJ, Johnson JE (2011) Ascl1 (Mash1) defines cells with long-term neurogenic potential in subgranular and subventricular zones in adult mouse brain. PLoS One 6:e18472.

Kinzler KW, Ruppert JM, Bigner SH, Vogelstein B (1988) The GLI gene is a member of the Kruppel family of zinc finger proteins. Nature 332:371–374.

Kramer PR, Wray S (2000) Midline nasal tissue influences nestin expression in nasal-placode-derived luteinizing hormone-releasing hormone neurons during development. Dev Biol 227:343–357.

Krauss S, Foerster J, Schneider R, Schweiger S (2008) Protein phosphatase 2A and rapamycin regulate the nuclear localization and activity of the transcription factor GLI3. Cancer Res 68:4658–4665.

Kroisel PM, Petek E, Wagner K (2001) Phenotype of five patients with Greig syndrome and microdeletion of 7p13. Am J Med Genet 102:243–249.

Lavdas AA, Grigoriou M, Pachnis V, Parnavelas JG (1999) The medial ganglionic eminence gives rise to a population of early neurons in the developing cerebral cortex. J Neurosci 19:7881–7888.

Leanos-Miranda A, Ulloa-Aguirre A, Janovick JA, Conn PM (2005) In vitro coexpression and pharmacological rescue of mutant gonadotropin-releasing hormone receptors causing hypogonadotropic hypogonadism in humans expressing compound heterozygous alleles. J Clin Endocrinol Metab 90:3001–3008.

Lewkowitz-Shpuntoff HM, Hughes VA, Plummer L, Au MG, Doty RL, Seminara SB, Chan YM, Pitteloud N, Crowley WF, Jr., Balasubramanian R Olfactory phenotypic spectrum in idiopathic hypogonadotropic hypogonadism: pathophysiological and genetic implications. J Clin Endocrinol Metab 97:E136–144.

Lin JM, Taroc EZM, Frias JA, Prasad A, Catizone AN, Sammons MA, Forni PE (2018a) The transcription factor Tfap2e/AP-2ε plays a pivotal role in maintaining the identity of basal vomeronasal sensory neurons. bioRxiv.

Lin JM, Taroc EZM, Frias JA, Prasad A, Catizone AN, Sammons MA, Forni PE (2018b) The transcription factor Tfap2e/AP-2epsilon plays a pivotal role in maintaining the identity of basal vomeronasal sensory neurons. Dev Biol.

Markram H, Toledo-Rodriguez M, Wang Y, Gupta A, Silberberg G, Wu C (2004) Interneurons of the neocortical inhibitory system. Nat Rev Neurosci 5:793–807.

Matera I, Watkins-Chow DE, Loftus SK, Hou L, Incao A, Silver DL, Rivas C, Elliott EC, Baxter LL, Pavan WJ (2008) A sensitized mutagenesis screen identifies Gli3 as a modifier of Sox10 neurocristopathy. Hum Mol Genet 17:2118–2131.

Mayeur A, Duclos C, Honore A, Gauberti M, Drouot L, do Rego JC, Bon-Mardion N, Jean L, Verin E, Emery E, Lemarchant S, Vivien D, Boyer O, Marie JP, Guerout N (2013) Potential of olfactory ensheathing cells from different sources for spinal cord repair. PLoS One 8:e62860.

McNay DE, Pelling M, Claxton S, Guillemot F, Ang SL (2006) Mash1 is required for generic and subtype differentiation of hypothalamic neuroendocrine cells. Mol Endocrinol 20:1623–1632.

Metz H, Wray S (2010) Use of mutant mouse lines to investigate origin of gonadotropin-releasing hormone-1 neurons: lineage independent of the adenohypophysis. Endocrinology 151:766–773.

Miller AM, Treloar HB, Greer CA (2010) Composition of the migratory mass during development of the olfactory nerve. J Comp Neurol 518:4825–4841.

Miller SR, Perera SN, Benito C, Stott SR, Baker CV (2016) Evidence for a Notch1-mediated transition during olfactory ensheathing cell development. J Anat 229:369–383.

Miller SR, Benito C, Mirsky R, Jessen KR, Baker CVH (2018) Neural crest Notch/Rbpj signaling regulates olfactory gliogenesis and neuronal migration. Genesis 56:e23215.

Naruse I, Fukui Y, Keino H, Taniguchi M (1994) The arrest of luteinizing hormone-releasing hormone neuronal migration in the genetic arhinencephalic mouse embryo (Pdn/Pdn). Brain Res Dev Brain Res 81:178–184.

Niewiadomski P, Kong JH, Ahrends R, Ma Y, Humke EW, Khan S, Teruel MN, Novitch BG, Rohatgi R (2014) Gli protein activity is controlled by multisite phosphorylation in vertebrate Hedgehog signaling. Cell Rep 6:168–181.

Ohtsuka T, Sakamoto M, Guillemot F, Kageyama R (2001) Roles of the basic helix-loop-helix genes Hes1 and Hes5 in expansion of neural stem cells of the developing brain. J Biol Chem 276:30467–30474.

Ohtsuka T, Ishibashi M, Gradwohl G, Nakanishi S, Guillemot F, Kageyama R (1999) Hes1 and Hes5 as notch effectors in mammalian neuronal differentiation. EMBO J 18:2196–2207.

Pablo Mendez J, Zenteno JC, Coronel A, Soriano-Ursua MA, Valencia-Villalvazo EY, Soderlund D, Coral-Vazquez RM, Canto P (2014) Triallelic digenic mutation in the prokineticin 2 and GNRH receptor genes in two brothers with normosmic congenital hypogonadotropic hypogonadism. Endocr Res:1–6.

Pilaz LJ, Patti D, Marcy G, Ollier E, Pfister S, Douglas RJ, Betizeau M, Gautier E, Cortay V, Doerflinger N, Kennedy H, Dehay C (2009) Forced G1-phase reduction alters mode of division, neuron number, and laminar phenotype in the cerebral cortex. Proc Natl Acad Sci U S A 106:21924–21929.

Pingault V, Bodereau V, Baral V, Marcos S, Watanabe Y, Chaoui A, Fouveaut C, Leroy C, Verier-Mine O, Francannet C, Dupin-Deguine D, Archambeaud F, Kurtz FJ, Young J, Bertherat J, Marlin S, Goossens M, Hardelin JP, Dode C, Bondurand N (2013) Loss-of-function mutations in SOX10 cause Kallmann syndrome with deafness. Am J Hum Genet 92:707–724.

Pitteloud N, Hayes FJ, Dwyer A, Boepple PA, Lee H, Crowley WF, Jr. (2002) Predictors of outcome of long-term GnRH therapy in men with idiopathic hypogonadotropic hypogonadism. J Clin Endocrinol Metab 87:4128–4136.

Pitteloud N et al. (2007) Digenic mutations account for variable phenotypes in idiopathic hypogonadotropic hypogonadism. J Clin Invest 117:457–463.

Prince JE, Cho JH, Dumontier E, Andrews W, Cutforth T, Tessier-Lavigne M, Parnavelas J, Cloutier JF (2009) Robo-2 controls the segregation of a portion of basal vomeronasal sensory neuron axons to the posterior region of the accessory olfactory bulb. J Neurosci 29:14211–14222.

Quaynor SD, Kim HG, Cappello EM, Williams T, Chorich LP, Bick DP, Sherins RJ, Layman LC (2011) The prevalence of digenic mutations in patients with normosmic hypogonadotropic hypogonadism and Kallmann syndrome. Fertil Steril 96:1424–1430 e1426.

Quaynor SD, Bosley ME, Duckworth CG, Porter KR, Kim SH, Kim HG, Chorich LP, Sullivan ME, Choi JH, Cameron RS, Layman LC (2016) Targeted next generation sequencing approach identifies eighteen new candidate genes in normosmic hypogonadotropic hypogonadism and Kallmann syndrome. Mol Cell Endocrinol 437:86–96.

Rich CA, Perera SN, Andratschke J, Stolt CC, Buehler DP, Southard-Smith EM, Wegner M, Britsch S, Baker CVH (2018) Olfactory ensheathing cells abutting the embryonic olfactory bulb express Frzb, whose deletion disrupts olfactory axon targeting. Glia 66:2617–2631.

Richter MW, Fletcher PA, Liu J, Tetzlaff W, Roskams AJ (2005) Lamina propria and olfactory bulb ensheathing cells exhibit differential integration and migration and promote differential axon sprouting in the lesioned spinal cord. J Neurosci 25:10700–10711.

Saitsu H, Komada M, Suzuki M, Nakayama R, Motoyama J, Shiota K, Ishibashi M (2005) Expression of the mouse Fgf15 gene is directly initiated by Sonic hedgehog signaling in the diencephalon and midbrain. Dev Dyn 232:282–292.

Sasaki H, Hui C, Nakafuku M, Kondoh H (1997) A binding site for Gli proteins is essential for HNF-3beta floor plate enhancer activity in transgenics and can respond to Shh in vitro. Development 124:1313–1322.

Sasaki H, Nishizaki Y, Hui C, Nakafuku M, Kondoh H (1999) Regulation of Gli2 and Gli3 activities by an amino-terminal repression domain: implication of Gli2 and Gli3 as primary mediators of Shh signaling. Development 126:3915–3924.

Schimmang T, Lemaistre M, Vortkamp A, Ruther U (1992) Expression of the zinc finger gene Gli3 is affected in the morphogenetic mouse mutant extra-toes (Xt). Development 116:799–804.

Schwanzel-Fukuda M, Pfaff DW (1989) Origin of luteinizing hormone-releasing hormone neurons. Nature 338:161–164.

Sykiotis GP, Plummer L, Hughes VA, Au M, Durrani S, Nayak-Young S, Dwyer AA, Quinton R, Hall JE, Gusella JF, Seminara SB, Crowley WF, Jr., Pitteloud N (2010) Oligogenic basis of isolated gonadotropin-releasing hormone deficiency. Proceedings of the National Academy of Sciences of the United States of America 107:15140–15144.

Taroc EZM, Prasad A, Lin JM, Forni PE (2017) The terminal nerve plays a prominent role in GnRH-1 neuronal migration independent from proper olfactory and vomeronasal connections to the olfactory bulbs. Biol Open 6:1552–1568.

Veistinen L, Takatalo M, Tanimoto Y, Kesper DA, Vortkamp A, Rice DP (2012) Loss-of-Function of Gli3 in Mice Causes Abnormal Frontal Bone Morphology and Premature Synostosis of the Interfrontal Suture. Front Physiol 3:121.

Wang H, Ge G, Uchida Y, Luu B, Ahn S (2011) Gli3 is required for maintenance and fate specification of cortical progenitors. J Neurosci 31:6440–6448.

Wichterle H, Garcia-Verdugo JM, Herrera DG, Alvarez-Buylla A (1999) Young neurons from medial ganglionic eminence disperse in adult and embryonic brain. Nat Neurosci 2:461–466.

Wichterle H, Turnbull DH, Nery S, Fishell G, Alvarez-Buylla A (2001) In utero fate mapping reveals distinct migratory pathways and fates of neurons born in the mammalian basal forebrain. Development 128:3759–3771.

Wray S, Grant P, Gainer H (1989) Evidence that cells expressing luteinizing hormone-releasing hormone mRNA in the mouse are derived from progenitor cells in the olfactory placode. Proceedings of the National Academy of Sciences of the United States of America 86:8132–8136.

Young J, Metay C, Bouligand J, Tou B, Francou B, Maione L, Tosca L, Sarfati J, Brioude F, Esteva B, Briand-Suleau A, Brisset S, Goossens M, Tachdjian G, Guiochon-Mantel A (2012) SEMA3A deletion in a family with Kallmann syndrome validates the role of semaphorin 3A in human puberty and olfactory system development. Hum Reprod 27:1460–1465.

Yu T, Fotaki V, Mason JO, Price DJ (2009) Analysis of early ventral telencephalic defects in mice lacking functional Gli3 protein. J Comp Neurol 512:613–627.

